# A New Monocyte Epigenetic Clock Reveals Effects of Alcohol Consumption on Epigenetic Aging in Three Independent Cohorts

**DOI:** 10.1101/2021.03.22.436488

**Authors:** Xiaoyu Liang, Rajita Sinha, Amy C. Justice, Mardge H. Cohen, Bradley E. Aouizerat, Ke Xu

## Abstract

**Background:** Excessive alcohol consumption increases the risk of aging-related comorbidities and mortality. Assessing the impact of alcohol consumption on biological age is important for clinical decision-making and prevention. Evidence shows that alcohol alters monocyte function, and age is associated with DNA methylome and transcriptomic changes among monocytes. However, no monocyte-based epigenetic clock is currently available. In this study, we developed a new monocyte-based DNA methylation clock (MonoDNAmAge) by using elastic net regularization. The MonoDNAmAge was validated by benchmarking using epigenetic age acceleration (EAA) in HIV infection. Using MonoDNAmAge clock as well as four established clocks (i.e., HorvathDNAmAge, HannumDNAmAge, PhenoDNAmAge, GrimDNAmAge), we then evaluated the effect of alcohol consumption on biological aging in three independent cohorts (N=2,242).

**Results:** MonoDNAmAge, comprised of 186 CpG sites, was highly correlated with chronological age (rtraining=0.96, p<2.20E-16; rtesting=0.86, p=1.55E-141). The MonoDNAmAge clock predicted an approximately 10-year age acceleration from HIV infection in two cohorts. Quadratic regression analysis showed a nonlinear relationship between MonoDNAmAge and alcohol consumption in the Yale Stress Center Community Study (YSCCS, *p*_*model*_=4.55E-08, *p*_*x*^2^_ =7.80E-08) and in the Veteran Aging Cohort Study (VACS, *p*_*model*_=1.85E-02, *p*_*x*^2^_ =3.46E-02). MonoDNAmAge and light alcohol consumption showed a negative linear relationship in the Women’s Interagency HIV Study (WIHS, *β*=-2.63, *p*_*x*_=2.82E-06). Heavy consumption increased EAAMonoDNAmAge up to 1.60 years in the VACS while light consumption decreased EAAMonoDNAmAge to 2.66 years in the WIHS. These results were corroborated by the four established epigenetic clocks.

**Conclusions:** We observed a nonlinear effect of alcohol consumption on epigenetic age that is estimated by a novel monocyte-based “clock” in three distinct cohorts, highlighting the complex effects of alcohol consumption on biological age.

## Background

Alcohol consumption has significant adverse effects on health and contributes to increased morbidity and mortality [1]. Biological aging has been proposed as an indicator of alcohol’s adverse effect on health and frailty [2, 3]. Recently developed epigenetic “clocks” employ cellular DNA methylation (DNAm) as a measure of the aging process. DNAm-based epigenetic clocks were shown to be more sensitive and precise measures of cellular age than other genomic measures (e.g., transcriptome, telomere length) [4]. However, the study of the impact of alcohol consumption on epigenetic age is still in its infancy.

To date, more than a dozen DNAm-based epigenetic clocks have been reported including four well-established DNAm-based age estimators. The Horvath clock (HorvathDNAmAge) is based on 353 CpG sites capture multi-tissue biological age estimation [5]. The Hannum clock (HannumDNAmAge) is derived from 71 CpGs in leukocytes [6]. The Levine clock (PhenoDNAmAge) employs 513 CpGs predicting life span [7]. Lu’s GrimAge clock (GrimDNAmAge) is a linear combination of chronological age, sex, and 1,030 CpG sites modeled as surrogate biomarkers for seven plasma proteins and smoking pack-years, predicting age at death [8]. These clocks have been applied as biomarkers of the aging process to understand how numerous medical or psychiatric conditions impact biological age. For example, HIV infection has shown a 5 to 10 year accelerated biological age [9, 10].

These established “clocks” have been applied to examine alcohol use on biological age. Epigenetic age acceleration (EAA) in heavy alcohol use and in children with fetal alcohol spectrum disorder have been reported [11, 12]. People with alcohol use disorder show a trend in age acceleration in liver tissue [13] and have a 2.22-year acceleration in blood relative to healthy individuals [14]. On the other hand, light to moderate alcohol use appears to no change or even slow extrinsic epigenetic age acceleration [15, 16].

These observations suggest that the impact of alcohol consumption on the biological aging process may differ by the quantity of use and by tissue types.

One limitation of previous studies using available epigenetic clocks is the lack of tissue and cell-type specificity to estimate the biological age of alcohol use, thus they may offer limited insight into the mechanisms of tissue-specific cellular aging. Alcohol consumption changes immunity and inflammatory functions that may result from functional alteration of immune cells [17]. For example, moderate or heavy alcohol use changes monocyte function [18–20]. Interestingly, Szabo et al [19] reported that alcohol consumption had a biphasic effect on interferon-inducibility and could affect monocyte-derived inflammatory cytokine production, which changes the course of the aging process. In addition, monocytes show aging-related gene dysfunction in metabolism, immune function, and inflammation [21, 22], as well as DNAm and transcriptomic alterations [23, 24]. Therefore, it is reasonable to posit that a set of CpG sites selected from monocytes can serve as an indicator of the impact of alcohol consumption on biological age and provide insight into its underlying mechanisms.

In this study, we aimed to characterize the effect of a range of alcohol consumption on monocyte epigenetic age. We developed a novel “clock” derived from the human monocyte DNA methylome (termed “MonoDNAmAge”) by using Elastic Net Regularization (ENR). We evaluated the performance of the MonoDNAmAge in estimating HIV-associated age acceleration as a benchmark. Finally, we assessed the impact of alcohol consumption on MonoDNAmAge, EAA, and apparent methylation age rate (AMAR) in three distinct cohorts: Yale Stress Center Community Cohort (YSCCS) [25]; Veteran Aging Study Cohort (VACS) [26]; and Women’s Interagency HIV Study (WIHS) [27, 28]. Demographic and phenotypic information for each cohort is presented in **Table 1 and Supplementary Information**. Four well-established clocks (i.e., HorvathDNAmAge, HannumDNAmAge, PhenoDNAmAge, GrimDNAmAge) were also used to assess the impact of alcohol consumption on measures of biological age (**Fig. 1**).

**Fig. 1.**
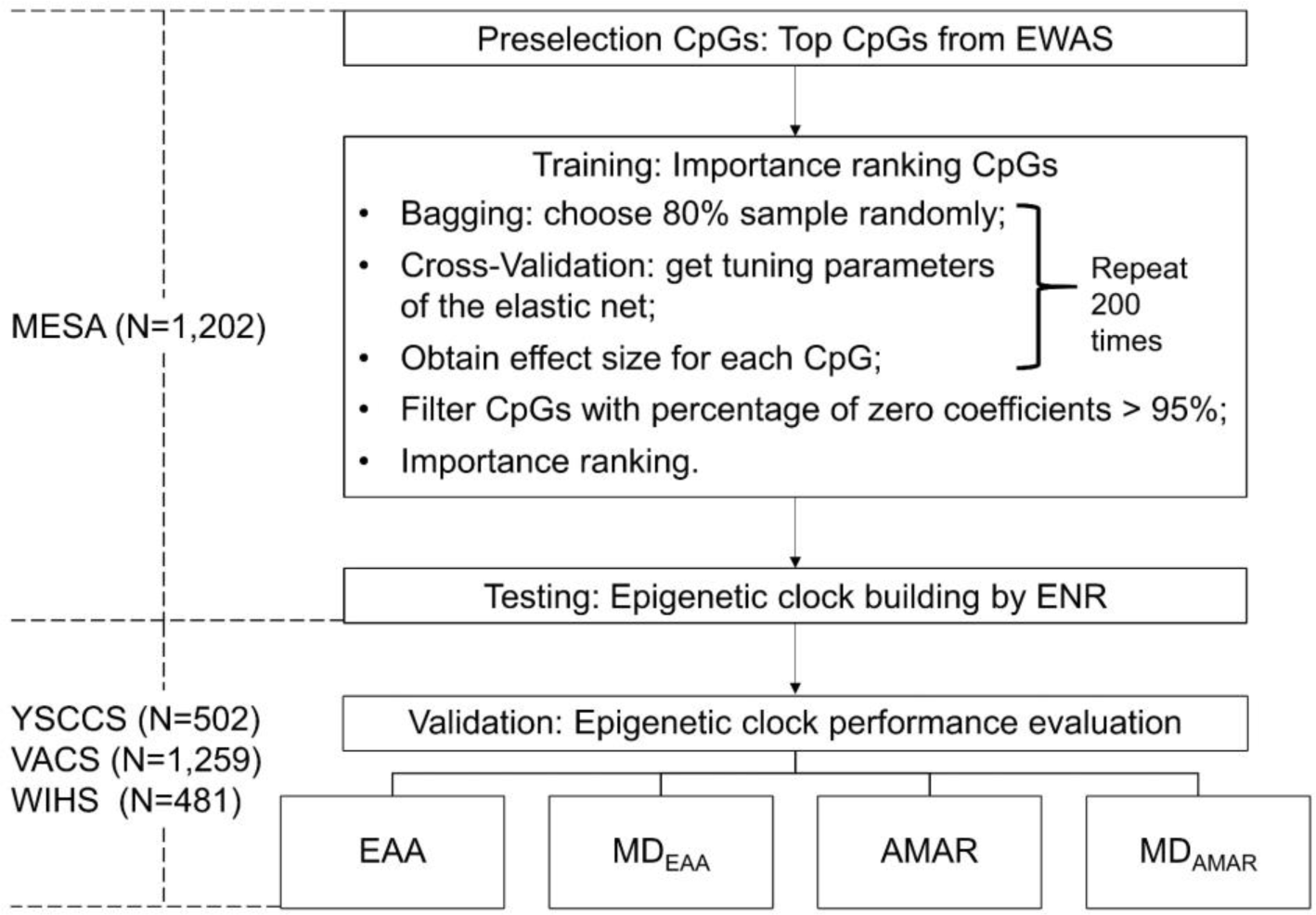
The workflow of establishing a monocyte DNA methylation “clock” using a feature selection method. YSCCS: Yale Stress Center Cohort Study; VACS: Veterans Aging Cohort Study; WIHS: Women’s Interagency HIV Study; EWAS: Epigenome-Wide Association Studies; ENR: Elastic Net Regularization; EAA: Epigenetic Age Acceleration, the residuals of regressing DNA methylation age on chronological age; MDEAA: The mean difference of EAA between different groups for phenotypes of interest; AMAR: Apparent Methylation Age Rate, the ratio of DNA methylated age to chronological age; MDAMAR: The mean difference of AMAR between different groups for phenotypes of interest.

**Table 1.** Demographic and clinical characteristics for YSCCS, VACS, and WIHS

| Variable | YSCCS |  |  | VACS |  |  | WIHS |  |  |
| --- | --- | --- | --- | --- | --- | --- | --- | --- | --- |
|  | HAD | MAD | p | HAD | MAD | p | LAD | NAD | p |
|  | Men:<br>AUDIT≥8<br>Women:<br>AUDIT≥7<br>(N=148) | Men:<br>AUDIT<8<br>Women:<br>AUDIT<7<br>(N=354) |  | PEth≥20<br>(N=299) | PEth<20<br>(N=738) |  | 0<NDRNKWK≤7<br>(N=196) | NDRNKWK=0<br>(N=255) |  |
| age | **26.8 ± 7.13** | **29.8 ± 9.28** | **1.35E-04** | 50.7 ± 7.23 | 51.1 ± 8.14 | 5.03E-01 | **40.3 ± 9.13** | **43.7 ± 9.46** | **1.13E-04** |
| sex (%male) | **64.9%** | **35.3%** | **2.18E-09** | 100% | 100%/ | NA | 0% | 0% | NA |
| race (AA) | **12.2%** | **22.4%** | **1.29E-02** | **86.6%** | **80.2%** | **1.90E-02** | 49.0% | 49.8% | 9.69E-01 |
| smoker | **39.9%** | **12.7%** | **1.77E-11** | **68.1%** | **54.3%** | **6.27E-05** | 70.9% | 67.3% | 8.97E-01 |
| HIV-positive | 0% | 0% | NA | 93.7% | 91.3% | 2.63E-01 | **50.5%** | **63.9%** | **1.54E-04** |
| log<sub>10</sub>VL | 0 | 0 | NA | 2.82 ± 1.23 | 2.66 ± 1.24 | 8.44E-02 | 2.16 ± 0.69 | 2.05 ± 0.48 | 1.34E-01 |
| cannabis use | **88.4%** | **56.9%** | **2.40E-11** | **84.5%** | **73.4%** | **2.21E-04** | **50.8%** | **29.1%** | **3.83E-06** |
| cocaine use | NA | NA | NA | **75.3%** | **66.0%** | **4.34E-03** | NA | NA | NA |
| stimulant use | NA | NA | NA | 36.9% | 39.9% | 4.08E-01 | NA | NA | NA |
| opiate use | NA | NA | NA | 42.3% | 44.2% | 6.27E-01 | NA | NA | NA |

YSCCS: Yale Stress Center Cohort Study; VACS: Veterans Aging Cohort Study; WIHS: Women's Interagency HIV Study.
HAD: Heavy Alcohol Drinking; MAD: Moderate Alcohol Drinking; LAD: Light Alcohol Drinking; NAD: Non-Alcohol Drinking.
AA: African American, AUDIT: Alcohol Use Disorders Identification Test, AUDIT-C: Alcohol Use Disorders Identification Test-Consumption, VL: viral load.
Welch's two-sample t-test for testing two groups mean; chi-square test for testing two groups percentage.
Significant phenotypes are in bold.
The cannabis, cocaine, stimulant, and opiate use in VACS were defined as case (>0) and control (=0).

## Results

### MonoDNAmAge clock and biological interpretation

MonoDNAmAge was derived from CD14+ monocyte methylome in Multi-Ethnic Study of Atherosclerosis (MESA) (N=1,202) (GSE56046) [24]. We selected a set of 186 age-associated CpGs using ENR. The correlation between DNAm age and chronological age was 0.96 (p<2.20E-16) in the training set and was 0.86 (p=1.55E-141) in the testing set (**Fig. 2a**). These 186 CpG sites mapped to 135 genes (**Supplementary Information: Table S1**), including well-established genes associated with age (e.g. *KLF14*), transcription factors (e.g. *RUNX3*), and inflammatory function (e.g. *IL17RC*).

**Fig. 2.**
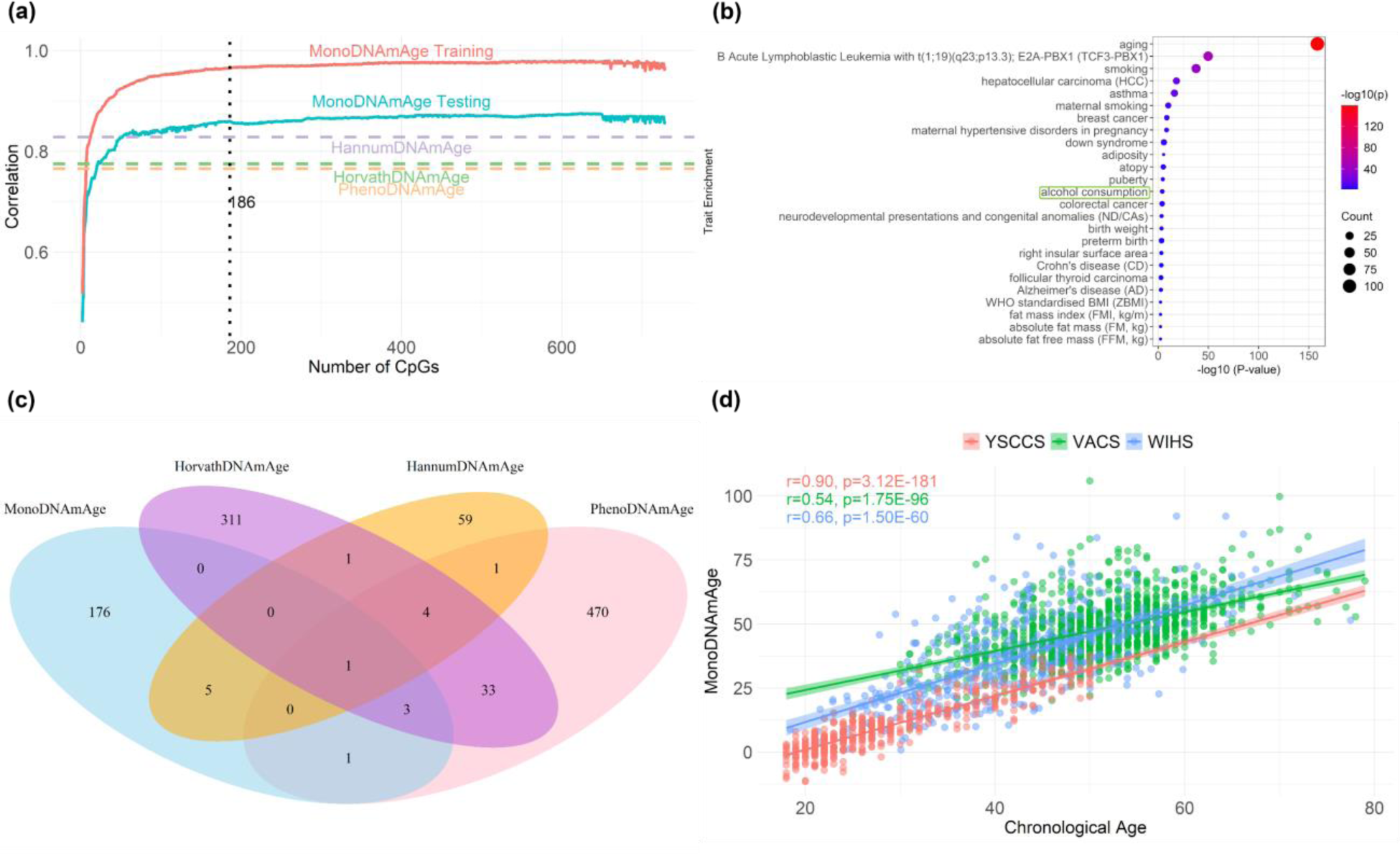
MonoDNAmAge development and performance. (a) Feature selection using elastic net regularization (ENR) for estimating MonoDNAmAge. The figure shows the Pearson correlation coefficients for the training and the testing data sets from MESA. **(b)** Trait enrichment of the selected 186 CpGs for MonoDNAmAge. **(c)** Overlapping CpG sites among four epigenetic clocks. **(d)** Correlation between MonoDNAmAge and chronological age in three independent cohorts: Yale Stress Center Cohort Study (YSCCS); Veterans Aging Cohort Study (VACS); and Women’s Interagency HIV Study (WIHS).

Interestingly, the 186 CpG sites were enriched for 25 complex traits with p<5.00E-03 in the Epigenome-Wide Association Study (EWAS) Atlas database [29] (**Fig. 2b**). The top significant traits included aging (p=2.57E-159), smoking (p=2.39E-38), breast cancer (p=2.71E-09), and alcohol consumption (p=8.95E-05). Three established clocks (HorvathDNAmAge, HannumDNAmAge, and PhenoDNAmAge) were enriched for multiple traits including age and smoking, but except HorvathDNAmAge (p=2.45E-03), none of the other two clocks was enriched for alcohol consumption (**Supplementary Information: Fig. S1**). One CpG site, *SCGN* cg06493394 was shared across the four clocks (**Fig. 2c**). Identifiers of CpG sites for GrimDNAmAge were not publicly available for comparisons. The 135 genes harboring the 186 CpG sites were enriched on biological pathways relevant to aging (e.g. regulation of multicellular organismal process) by performing Database for Annotation, Visualization and Integrated Discovery (DAVID) pathway enrichment analysis (http://david.niaid.nih.gov) (**Supplementary Information: Fig. S2**) [30].

MonoDNAmAge was significantly correlated with chronological age in all three cohorts (YSCCS: *r*=0.90, p=3.12E-181; VACS: *r*=0.54, p=1.75E-96; WIHS: *r*=0.66, p=1.50E-60) (**Fig. 2d**). We also estimated the correlations of four established clocks with chronological age in these three cohorts. As expected, all clocks showed significant correlations with chronological age in each cohort (YSCCS: p=8.42E-215∼4.69E-168; VACS: p=6.82E-205∼3.94E-135; WIHS: p=2.25E-175∼1.75E-91) (**Supplementary Information: Fig. S3**).

### Benchmarking MonoDNAmAge against HIV infection shows epigenetic age acceleration

EAA analysis showed an average age acceleration of 10.14-years in the VACS (pVACS=1.17E-24) and 12.17-years (pWIHS=2.07E-28) in the WIHS in HIV+ participants compared to HIV-negative (HIV-) participants (**Fig. 3a, Supplementary Information: Table S2**). To avoid a bias due to different sample sizes of HIV+ and HIV-participants, we randomly selected HIV+ participants to match the number of HIV-participants and compared the EAA value between the groups in both cohorts in **Fig. 3b**. As shown in **Fig. 3b**, the majority of HIV+ participants displayed positive EAA values while the majority of HIV-participants displayed negative EAA values. The methylation age rate (i.e., AMAR) of monocyte biological age was significantly greater in HIV-positive (HIV+) than HIV-participants in both cohorts (MDVACS=0.21, pVACS=1.00E-27; MDWIHS=0.34, pWIHS=6.51E-37) (**Fig. 3c, Supplementary Information: Table S2**).

**Fig. 3.**
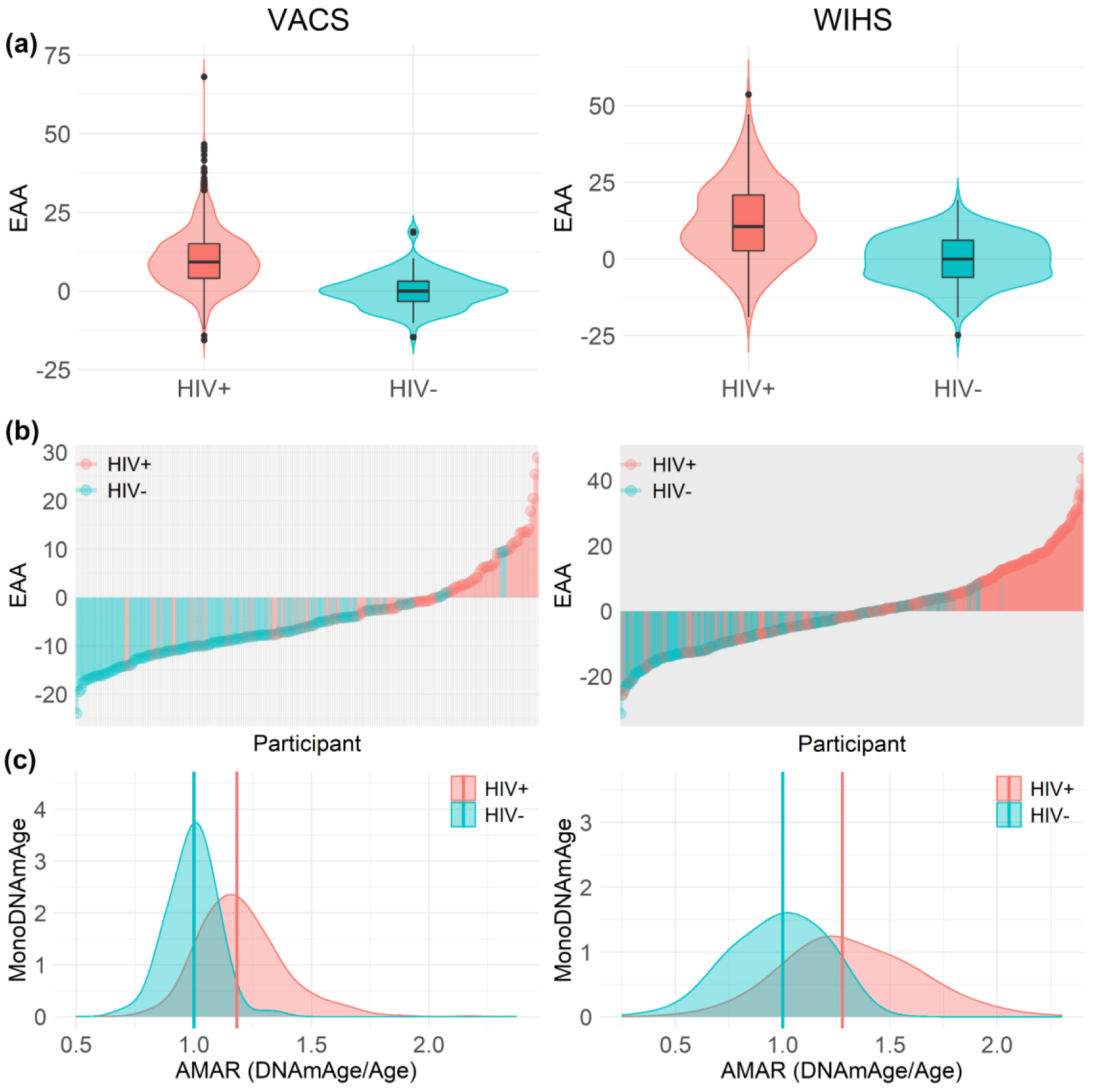
Comparison of MonoDNAmAge-based estimations of biological aging for HIV-positive and HIV-negative participants. (a) Violin plots showing significant differences in the Epigenetic Age Acceleration (EAA, the residuals of regressing MonoDNAmAge on chronological age) between HIV-positive (HIV+) and HIV-negative (HIV-) groups. **(b)** Lollipop plots for EAA between HIV+ and HIV-. **(c)** Density plots for Apparent Methylation Age Rate (AMAR, the ratio of MonoDNAmAge to chronological age) between HIV+ and HIV-. VACS: Veterans Aging Cohort Study; WIHS: Women’s Interagency HIV Study.

**Table 2.** Association between EAA/AMAR and alcohol consumption

| Measure | Cohort | Method | beta | 95% CI | p | MD |
| --- | --- | --- | --- | --- | --- | --- |
| EAA | YSCCS | MonoDNAmAge | 0.040 | (-0.031, 0.112) | 2.68E-01 | 0.129 |
|  |  | HorvathDNAmAge | 0.044 | (-0.008, 0.096) | 1.00E-01 | 0.415 |
|  |  | HannumDNAmAge | 0.028 | (-0.026, 0.083) | 3.11E-01 | 0.786 |
|  |  | PhenoDNAmAge | -0.031 | (-0.111, 0.049) | 4.42E-01 | -0.856 |
|  |  | GrimDNAmAge | 0.033 | (-0.013, 0.079) | 1.57E-01 | 0.767 |
|  | VACS | MonoDNAmAge | 0.477 | (0.153, 0.802) | **3.99E-03** | 1.603 |
|  |  | HorvathDNAmAge | 0.056 | (-0.141, 0.253) | 5.77E-01 | 0.135 |
|  |  | HannumDNAmAge | 0.237 | (0.038, 0.436) | **2.00E-02** | 0.711 |
|  |  | PhenoDNAmAge | 0.375 | (0.122, 0.628) | **3.76E-03** | 1.632 |
|  |  | GrimDNAmAge | 0.138 | (-0.079, 0.356) | 2.13E-01 | 1.520 |
|  | WIHS | MonoDNAmAge | -1.627 | (-2.463, -0.791) | **1.56E-04** | -2.656 |
|  |  | HorvathDNAmAge | -0.704 | (-1.054, -0.353) | **9.73E-05** | -0.806 |
|  |  | HannumDNAmAge | -0.676 | (-1.115, -0.236) | **2.75E-03** | -1.084 |
|  |  | PhenoDNAmAge | -0.843 | (-1.374, -0.312) | **1.98E-03** | -0.914 |
|  |  | GrimDNAmAge | -0.406 | (-0.658, -0.153) | **1.74E-03** | -0.746 |
| AMAR | YSCCS | MonoDNAmAge | -0.003 | (-0.0064, 0.0012) | 1.84E-01 | -0.056 |
|  |  | HorvathDNAmAge | 0.003 | (0.0007, 0.0044) | **7.29E-03** | 0.026 |
|  |  | HannumDNAmAge | 0.004 | (0.0021, 0.0065) | **1.33E-04** | 0.051 |
|  |  | PhenoDNAmAge | -0.003 | (-0.0058, 0.0001) | 6.32E-02 | -0.052 |
|  |  | GrimDNAmAge | 0.005 | (0.0025, 0.0070) | **3.31E-05** | 0.057 |
|  | VACS | MonoDNAmAge | 0.012 | (0.0054, 0.0185) | **3.55E-04** | 0.034 |
|  |  | HorvathDNAmAge | 0.002 | (-0.0026, 0.0063) | 4.19E-01 | 0.003 |
|  |  | HannumDNAmAge | 0.004 | (0, 0.0088) | **4.85E-02** | 0.012 |
|  |  | PhenoDNAmAge | 0.011 | (0.0054, 0.0161) | **9.16E-05** | 0.031 |
|  |  | GrimDNAmAge | 0.010 | (0.0044, 0.0157) | **4.89E-04** | 0.030 |
|  | WIHS | MonoDNAmAge | -0.046 | (-0.0668, -0.0247) | **2.54E-05** | -0.092 |
|  |  | HorvathDNAmAge | -0.015 | (-0.0241, -0.0062) | **1.00E-03** | -0.013 |
|  |  | HannumDNAmAge | -0.012 | (-0.0234, -0.0004) | **4.35E-02** | -0.009 |
|  |  | PhenoDNAmAge | -0.028 | (-0.0426, -0.013) | **2.61E-04** | -0.057 |
|  |  | GrimDNAmAge | -0.002 | (-0.0099, 0.0052) | 5.43E-01 | 0.002 |

EAA: Epigenetic Age Acceleration; AMAR: Apparent Methylation Age Rate.
YSCCS: Yale Stress Center Cohort Study; VACS: Veterans Aging Cohort Study; WIHS: Women's Interagency HIV Study.
95% CI: 95% confidence interval for coefficient beta.
The p-values in bold indicate the values less than 5.00E-02.

Additionally, we tested age acceleration in HIV infection using four established epigenetic clocks in VACS and WIHS. The EAAs estimated by all four clocks were significantly different between HIV+ and HIV-participants in both VACS (p=3.26E-11∼1.07E-02) and WIHS (p=4.62E-14∼1.34E-04). For example, the EAA differences between HIV+ and HIV-were as great as 2.53 in VACS and 3.60 in WIHS measured by HorvathDNAmAge (**Supplementary Information: Fig. S4 and Table S2**). AMAR was also greater in HIV+ than HIV-participants except for HannumDNAmAge in VACS and GrimDNAmAge in VACS and WIHS (**Supplementary Information: Fig. S5 and Table S2**).

### Nonlinear effects of alcohol consumption on DNA methylation age

ANOVA model comparison showed that the quadratic regression model fits the data better than the linear regression model for all five clocks in the YSCCS (p=6.27E-09∼2.41E-06) and VACS (p=8.32E-04∼3.91E-02) but not in the WIHS cohort (p=2.47E-01∼4.38E-01) (**Supplementary Information: Table S3**). Then, the quadratic regression was applied to YSCCS (**Fig. 4a**) and VACS (**Fig. 4b**). In the YSCCS cohort, we observed a nonlinear relationship between MonoDNAmAge and Alcohol Use Diagnosis Identification Test (AUDIT) score (*p*_*model*_ =4.55E-08, *p*_*x*^2^_ =7.80E-08) (**Fig. 4a**). The other four DNAm clocks also showed nonlinear associations between DNAm age and AUDIT score (HorvathDNAmAge: *p*_*model*_=2.82E-07, *p*_*x*^2^_ =2.37E-07; HannumDNAmAge: *p*_*model*_=5.54E-06, *p*_*x*^2^_ =2.41E-06; PhenoDNAmAge: *p*_*model*_=4.08E-10, *p*_*x*^2^_ =6.27E-09; GrimDNAmAge: *p*_*model*_=1.80E-07, *p*_*x*^2^_ =6.12E-08).

**Fig. 4.**
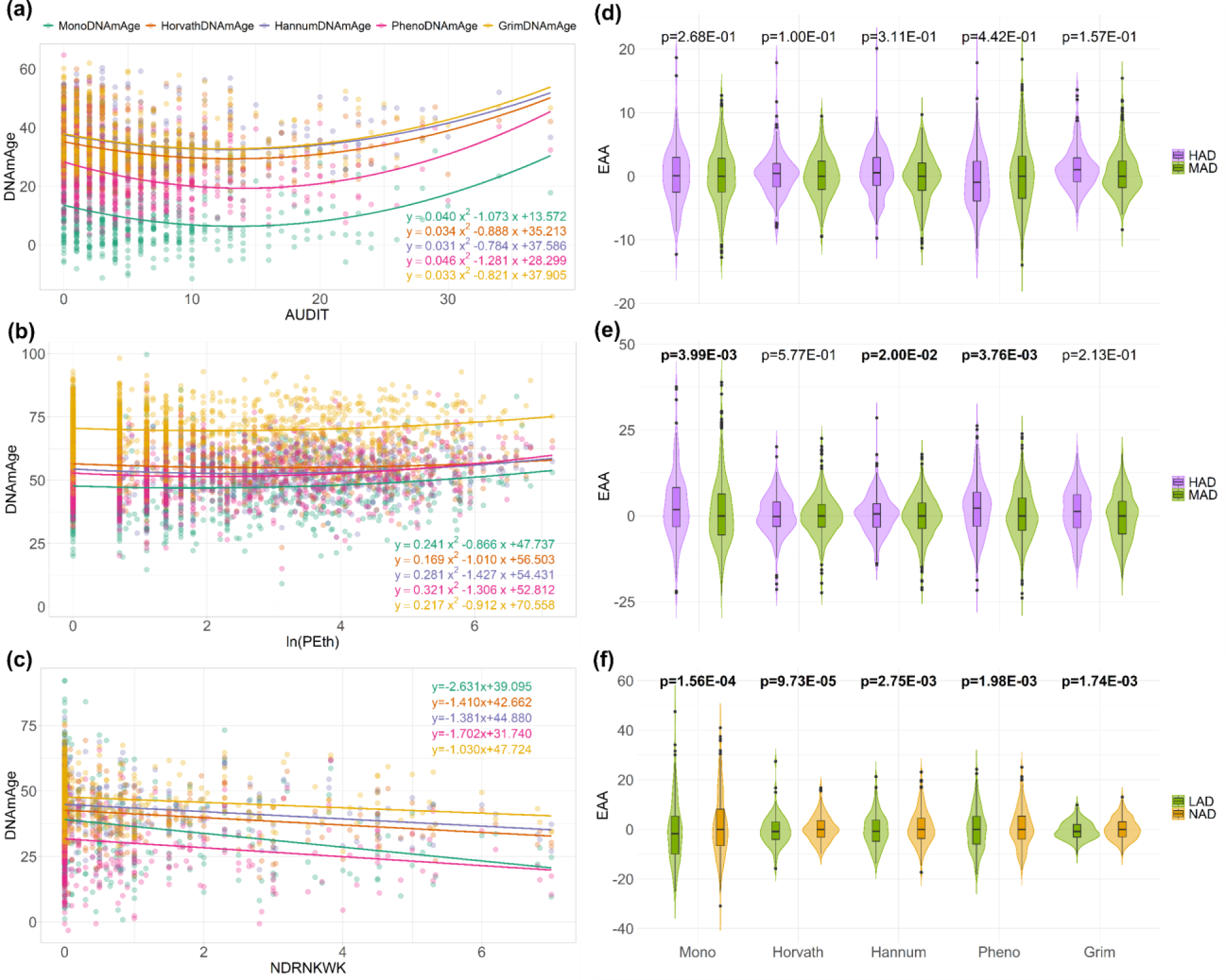
Relationships between DNA methylation age (DNAmAge) based on different approaches and alcohol consumption. The three rows correspond to Yale Stress Center Cohort Study (YSCCS), Veterans Aging Cohort Study (VACS), and Women’s Interagency HIV Study (WIHS). **(a)** The parabola shows the relationship between DNAmAge and the Alcohol Use Disorders Identification Test (AUDIT) score in the YSCCS. **(b)** The parabola shows the relationship between DNAmAge and the natural logarithm of Phosphatidylethanol (ln(PEth)) in the VACS. **(c)** The linear regression line shows the negative relationship between DNAmAge and the number of drinks per week (NDRNKWK) in the WIHS. **(d)-(f)** Violin plots for Epigenetic Age Acceleration (EAA) (the residuals of regressing DNA methylation age on chronological age) for different groups of alcohol consumption in the three cohorts. **(d)** Comparison of heavy alcohol drinking (HAD) (AUDIT>=7 for men and AUDIT>=8 for women) and moderate alcohol drinking (MAD) (AUDIT<7 for men and AUDIT<8 for women) in the YSCCS. **(e)** Comparison of HAD (PEth>=20) and MAD (PEth<20) in the VACS. **(f)** Comparison of light alcohol drinking (LAD) (0<NDRNKWK<7) and non-alcohol drinking (NAD) (NDRNKWK=0) in the WIHS.

To further investigate the relationship between MonoDNAmAge and different levels of alcohol consumption, we performed two linear regression analyses at the inflection point of the nonlinear distribution at the AUDIT score of 13 in YSCCS, separately (**Supplementary Information: Fig. S6**). We found that MonoDNAmAge was negatively associated with consumption at AUDIT<13 (p=5.36E-06) and positively associated with heavy drinking of AUDIT>13 (p=3.70E-03). Every unit change in the AUDIT score<13 was associated with a 1.20-year decrease in MonoDNAmAge in the non-heavy alcohol drinking (non-HAD) group and every unit change in AUDIT score>13 was associated with a 0.36-year increase in MonoDNAmAge in the heavy alcohol drinking (HAD) group.

In the VACS cohort, quadratic regression also showed a significant association of MonoDNAmAge on alcohol consumption measured by the natural logarithm of phosphatidylethanol (In(PEth)) (*p*_*model*_=1.85E-02, *p*_*x*^2^_ =3.46E-02) (**Fig. 4b**). The results from the three established clocks provided estimates consistent to MonoDNAmAge: HannumDNAmAge (*p*_*model*_=3.74E-03, *p*_*x*^2^_ =8.32E-04), PhenoDNAmAge (*p*_*model*_=4.97E-04, *p*_*x*^2^_ =8.60E-04), GrimDNAmAge (*p*_*model*_=3.40E-02, *p*_*x*^2^_ =2.21E-02). Here, the inflection point for the curve of PEth levels was close to the PEth cutoff for the definition of HAD. We found that the slope below the PEth inflection point (non-HAD: PEth<20, that is ln(PEth)<2.996) showed no significant (p=6.79E-01) association with

MonoDNAmAge while the slope above the PEth inflection point (HAD: PEth≥20, that is In(PEth)≥2.996), showed a significant positive association with MonoDNAmAge (p=4.39E-02) (**Supplementary Information: Fig. S6**). Every unit change of In(PEth) above the inflection point was associated with a 1.31-year increase in DNAm age.

However, we found that no significant association of DNAm age on AUDIT-Consumption (AUDIT-C, first 3 items of AUDIT) score for all five clocks (*p*_*model*_=1.79E-01∼9.01E-01) (**Supplementary Information: Fig. S7**), suggesting that PEth is a more accurate measure of alcohol consumption on DNAm age than self-reported AUDIT-C.

In the WIHS, MonoDNAmAge was linearly associated with light alcohol consumption measured by the number of drinks per week (NDRNKWK) (*β*=-2.63, *p*_*model*_=2.82E-06, *p*_*x*_=2.82E-06) (**Fig. 4c**). The negative linear relationship between DNAm age and light alcohol consumption was also observed with the four established clocks (HorvathDNAmAge: *β*=-1.41, *p*_*model*_=1.33E-05, *p*_*x*_=1.33E-05; HannumDNAmAge: *β*=-1.38, *p*_*model*_=4.99E-05, *p*_*x*_=4.99E-05; PhenoDNAmAge: *β*=-1.70, *p*_*model*_=7.74E-05, *p*_*x*_=7.74E-05; GrimDNAmAge: *β*=-1.03, *p*_*model*_=1.23E-03, *p*_*x*_=1.23E-03).

### Alcohol consumption alters epigenetic age acceleration

In the YSCCS cohort, alcohol consumption was not associated with EAA estimated using MonoDNAmAge or the four established clocks (**Fig. 4d and Table 2**). However, AMAR from HorvathDNAmAge, HannumDNAmAge, and GrimDNAmAge showed greater DNAm age than chronological age and were associated with alcohol consumption (β=0.003∼0.005, p=3.31E-05∼7.29E-03).

In the VACS cohort, EAA analysis showed that alcohol consumption significantly accelerated MonoDNAmAge by 1.60 years (β=0.477, p=3.99E-03) (**Fig. 4e and Table 2**). EAA estimated by HannumDNAmAge and PhenoDNAmAge also showed significant acceleration (β=0.237, p=2.0E-02 and β=0.375, p=3.76E-03, respectively). AMAR estimated by MonoDNAmAge was associated with alcohol consumption (β=0.012, p=3.55E-04). Similarly, PhenoDNAmAge showed a similar association (β=0.011, p=9.16E-05). HannumDNAmAge and GrimDNAmAge were also significantly positively correlated with alcohol consumption.

In the WIHS cohort, MonoDNAmAge EAA was correlated with alcohol consumption in an inverse relationship (β=-1.627, p=1.56E-04, MDEAA=-2.656) (**Fig. 4f and Table 2**). All five clocks showed decelerations of DNAm age with an average MDEAA of -1.24 years.

AMAR for MonoDNAmAge was negatively correlated with alcohol consumption (MonoDNAmAge: β=-0.046, p=2.54E-05). Similarly, each AMAR for HorvathDNAmAge, HannumDNAmAge, and PhenoDNAmAge, was significantly negatively correlated with alcohol consumption.

DNAm-based age estimated by the MonoDNAmAge clock and four other clocks were significantly correlated with each of the six cell types in all three cohorts (r=0.54∼0.93, p=8.42E-215∼1.50E-60) (**Fig. 5**). Correlations were calculated for each of the five clocks with six estimated cell types in the HAD and moderate alcohol drinking (MAD) groups separately in YSCCS and VACS, and light alcohol drinking (LAD) and non-alcohol drinking (NAD) in WIHS. The correlation patterns were similar between the two alcohol use groups across the three cohorts, suggesting the effect of cell-type composition on the performance of the five biological age estimators does not differ significantly between HAD and MAD, or between LAD and NAD groups.

**Fig. 5.**
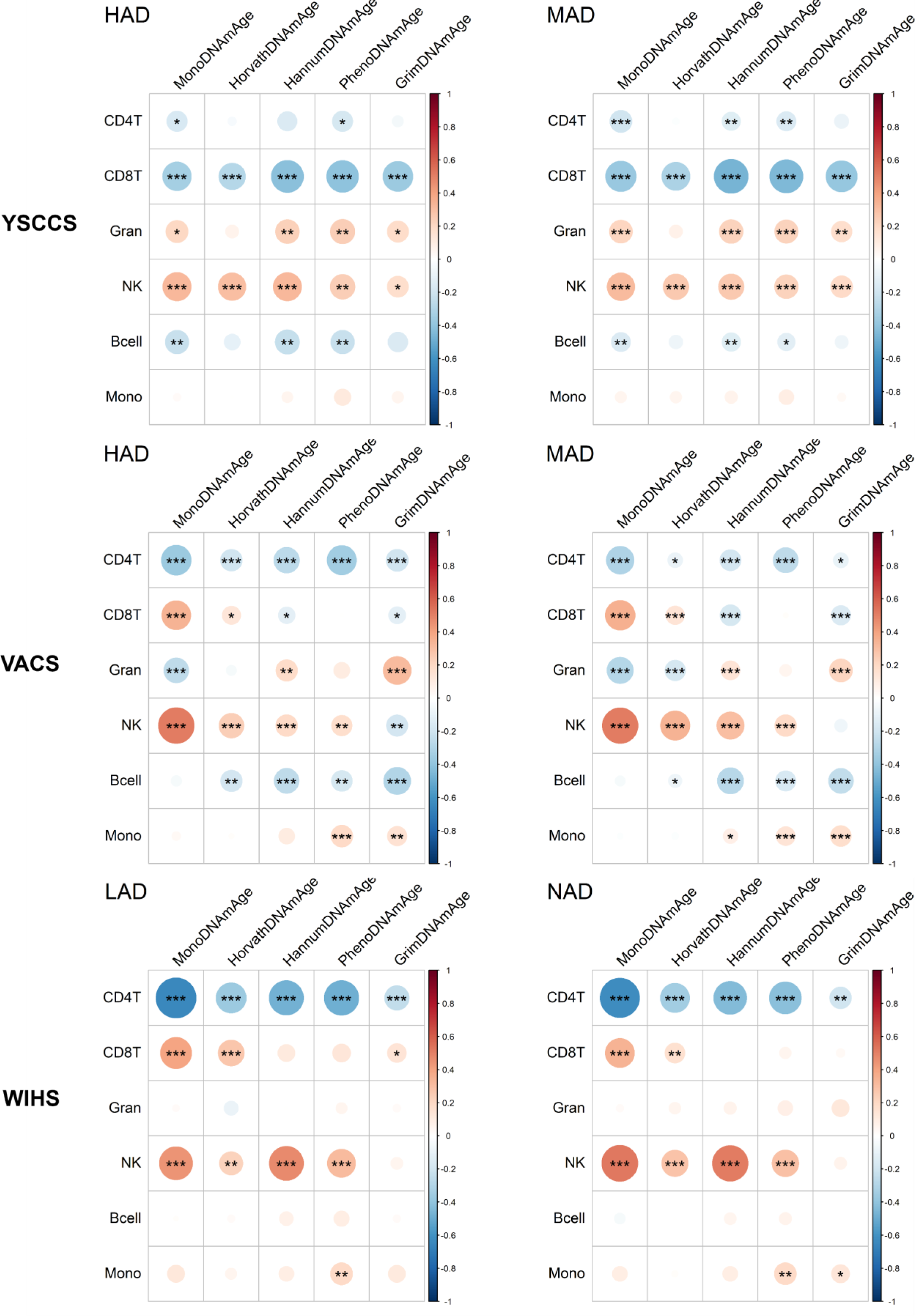
Correlation between five epigenetic clocks and cell-type proportions. Comparison of heavy alcohol drinkers (HAD) (AUDIT>=7 for men and AUDIT>=8 for women) and moderate alcohol drinkers (MAD) (AUDIT<7 for men and AUDIT<8 for women) in the Yale Stress Center Cohort Study (YSCCS); HAD (PEth>=20) and MAD (PEth<20) in the Veterans Aging Cohort Study (VACS); light alcohol drinkers (LAD) (0<NDRNKWK<7) and non-alcohol drinkers (NAD) (NDRNKWK=0) in the Women’s Interagency HIV Study (WIHS). ***indicates p-value<1.00E-03; ** indicates p-value<1.00E-02; * indicates p-value<5.00E-02.

## Discussion

Our results demonstrate that a set of DNAm CpG sites in monocytes are predictive of biological age, enabling the detection of the impact of alcohol consumption on DNAm age validated in three distinct cohorts. MonoDNAmAge clock, benchmarked against HIV infection, showed an approximate 10-year acceleration in HIV-positive participants.

More importantly, this novel MonoDNAmAge clock detected a nonlinear relationship between DNAm-based biological age and alcohol consumption in both a healthy community cohort (YSCCS) and a clinic-based cohort (VACS) that included heavy alcohol drinkers (∼33%). The impact of alcohol consumption on MonoDNAmAge was corroborated using four established epigenetic clocks. Thus, employing a comprehensive approach (i.e., one novel and four established epigenetic clocks, three independent cohorts, different but commonly employed measures of alcohol exposure), we identified for the first time that alcohol consumption appears to have a complex, nonlinear relationship with DNAm-based estimates of biological age.

Different from previous studies, here we examined the effect of alcohol consumption on epigenetic age by applying a novel cell-type-specific epigenetic clock related to the biological mechanism of alcohol consumption. The monocyte methylome plays an important role in epigenetic aging. Recently, differentially methylated regions in CD14 monocytes between young (24-30 years) and older (57-70 years) individuals have been reported [31]. Those age-associated CpG sites or DNAm regions have been linked to transcriptomic changes with aging [31]. Therefore, a CD14 monocyte epigenetic clock may provide a more accurate measure of the biological age of genes that compose the MonoDNAmAge clock that have functional implications for the aging process in monocyte-related phenotypes such as HIV infection and alcohol consumption. Indeed, the 186 MonoDNAmAge CpG sites are enriched in alcohol consumption and other related traits (e.g. smoking, cancer). Interestingly, although MonoDNAmAge and the established epigenetic clocks are composed of different sets of CpGs with the exception of one site (*SCGN* cg06493394), these clocks showed reasonably consistent but not identical performance in estimating alcohol’s effect on epigenetic age. We speculate that each epigenetic clock is estimating different but related facets of biological aging, inferred in part by the overlapping traits that show enrichment for the CpG sites that compose each DNAm-based epigenetic clock.

The nonlinear association between alcohol consumption and biological age is interesting. The beneficial and harmful effects of alcohol consumption have been well documented in the literature [1, 32–35]. While any amount of alcohol is harmful to health [1], some studies showed that alcohol consumption had a U-shaped relationship to biomarkers including HDL, LDL, VLDL, and inflammation [36, 37], underscoring the bidirectional effects on cardiovascular and metabolic disease risks. In a line of this evidence, we observed a nonlinear association between alcohol consumption and biological age in two cohorts, YSCCS and VACS, in which the numbers of heavy and non-heavy alcohol consumers were relatively balanced. In the WIHS cohort, the predominance of none-to-light drinkers may explain why the negative association with biological aging was similarly observed; insufficient observations were available to test the relationship between heavy drinkers and biological aging. The results of EAA showed heavy consumption measured by PEth accelerated epigenetic age of 1.60 years, consistent with a previous report of a 2-year acceleration in individuals with Alcohol Use Disorder (AUD) [14]. On the other hand, EAA was inversely associated with light to moderate alcohol consumption, which is consistent with a previous report that self-reported moderate alcohol use (i.e., number of drinks per week) showed a negative correlation with biological age estimated using PhenoDNAmAge [7].

A possible explanation for the observed nonlinear relationship between alcohol consumption and biological aging is that individuals with light to moderate alcohol use are more likely to follow a healthier lifestyle. For example, in the YSCCS and VACS, individuals not reporting HAD showed lower smoking rates and cannabis use, than those with HAD, suggesting that a healthy lifestyle may slow or potentially even reverse the epigenetic aging process due to alcohol consumption. Another possibility is due to underreported alcohol consumption among participants. For example, in the VACS cohort, 17.02% of participants reported AUDIT-C as 0 but PEth level among those participants are greater than 8ng/mL (**Supplementary Information: Fig. S8**), which indicates active alcohol use [38]. Inaccurately self-reported alcohol consumption may result in biased findings towards the slow acceleration of light-to-moderate drinkers on biological age. These observations need further investigations with accurately assessed phenotype or in longitudinal cohorts. Whereas the overall harm of alcohol drinking consumption is outweighed the benefit, causality has remained elusive [32].

We acknowledge some limitations of this study. A recent study suggests that the accuracy of DNAm-based epigenetic clock estimation is affected by sample size [39]. Validation of the MonoDNAmAge clock in a large independent sample is warranted. DNAm in the three cohorts was measured in DNA from whole blood. We expect that DNAm derived solely from monocytes may be a more accurate predictor of the impact of alcohol consumption. We are unable to examine DNAm age in past alcohol users, for different patterns of alcohol consumption, or for comorbidity with other substance or drug use. Furthermore, while differences in the effects of alcohol use between women and men are well documented, the study lacked sufficient power to evaluate for sex differences in alcohol exposure on biological aging.

### Conclusion

We found that alcohol use impacts epigenetic aging in a nonlinear fashion with heavy consumption accelerating while non-heavy use decreasing the acceleration of biological age. The use of cell-type-specific epigenetic clocks that are known to be directly impacted by alcohol exposure may provide more precisive information than provided by more holistic epigenetic clocks that estimate more global biological aging effects. Our study expands previous knowledge and provides new insights on the impact of a spectrum of alcohol use on epigenetic aging.

## Methods

### Study cohorts and phenotype assessments

***Multi-Ethnic Study of Atherosclerosis (MESA) (N=1,202)*** [24]. DNAm data of the MESA Epigenomics and Transcriptomics Study (GSE56046) was used to construct the MonoDNAmAge clock. The DNA samples were from CD14+ monocyte samples, collected from 1,202 individuals with ages ranging from 44 to 83 years.

The following three cohorts were used for assessing alcohol’s effects on DNAm aging (**Table 1,** see details of study cohorts and phenotype assessment in **Supplementary Information**).

***Yale Stress Center Cohort Study (YSCCS) (N=502)*** [25]. The cohort served as a community-based sample to examine the impact of alcohol consumption on DNAm age among healthy participants. The 10-item AUDIT scale was used to assess alcohol use. HAD was defined as AUDIT≥8 for men and AUDIT≥7 for women (N=148) [40]. The average AUDIT score was 5.69 among all participants in the cohort.

***Veterans Aging Cohort Study (VACS) (N=1,259)*** [26]. The cohort served as a clinic-based sample to benchmark HIV infection on MonoDNAmAge and to examine the effects of alcohol consumption on DNAm age. The participants included both HIV+ (N=1,151) and HIV-(N=104) individuals. A majority of HIV+ participants were on antiretroviral therapy and were virally suppressed (63.66%). Alcohol consumption was assessed by using PEth, a biomarker for alcohol use [22] that is positively correlated with AUDIT scores [41, 42]. HAD was defined as PEth≥20 (N=299) according to a previous study [43]. The average PEth was 41.7 ng/ml. Self-reported AUDIT-C was also collected for each participant.

***Women’s Interagency HIV Study (WIHS)* (N=481)** [27, 28]. The cohort served as a clinic-and community-based sample to examine HIV infection and alcohol consumption on DNAm age. This study included HIV-positive (N=272, 90.44% virally suppressed) and HIV-negative (N=209) participants. The cohort predominantly reported light alcohol use with an average NDRNKWK of 0.7. The LAD was defined as 0<NDRNKWK≤7 (N=196), and the NAD was defined as NDRNKWK=0 (N=255) [44].

### DNA methylation and data quality control (QC)

Epigenome-wide CpG methylation was profiled by using either the Illumina HumanMethylation450 BeadChip (HM450K) (San Diego, CA, USA) in MESA (monocyte), YSCCS (blood), and VACS (blood) (57.2% of the sample) or Illumina HumanMethylation EPIC BeadChip (EPIC) (San Diego, CA, USA) in 42.8% of the VACS samples and the WIHS (blood) samples. All samples in the three study cohorts (YSCCS, VACS, and WIHS) were processed at the Yale Center for Genomic Analysis [45]. We applied the method described by Houseman et al. [46, 47] to estimate proportions of CD4+ T cell, CD8+ T cell, NK T cell, B cell, monocyte, and granulocyte DNAm in each cohort. The QC for the MESA cohort was previously reported [24]. For the YSCCS, VACS, and WIHS cohorts, we applied the same QC process as reported in our previous studies [45, 48, 49].

### MonoDNAmAge clock development

As depicted in **Fig. 1**, we applied ENR to build a MonoDNAmAge clock. All samples in MESA were divided into two sets: a training (N=721) and a testing (N=481) set.

Briefly, the top 1,000 significant CpG features were preselected from monocyte methylome based on the EWAS on age in the MESA. In the training set, the ENR combined with random sampling was performed to select CpGs and establish the predictive model. The CpGs were filtered and sorted according to their coefficients in ENR. In the testing set, we added CpGs sequentially and calculated the corresponding correlations between the predicted MonoDNAmAge and chronological age. We determined the linear combination of the CpG set with the correlation at the inflection point of the performance curve in the testing set as MonoDNAmAge (**Fig. 2a**). The performance of MonoDNAmAge was evaluated in the three validation data sets (i.e., YSCCS, VACS, and WIHS). Analyses were performed using R software. ENR was performed using the function “cv.glmnet” in the “glmnet” package. The details of feature selection are described below.

#### Preselection of CpGs

Because a large number of CpGs may introduce noise, DNA methylation of CpGs under the epigenome-wide significance threshold may collectively account for phenotype variation and may improve prediction of a phenotype, we preselected the top 1000 age-associated CpGs from the EWAS on age in the MESA cohort.

#### Importance ranking of CpGs in the training set

The preselected CpGs were used to establish the predictive model in the training set of MESA. We randomly selected 80% of training subjects without replacement 200 times and constructed a model for each replication. We only included the CpGs that were present in more than 95% of all replications in the final model and ranked the CpGs based on the coefficients of all replications. For the ENR method in each replication, the 10-fold cross-validation procedure was included in the feature selection to obtain the estimations of ENR tuning parameters. We extracted the coefficients for the model with the lambda value corresponding to the minimum mean cross-validated error. The CpG importance ranking was based on the summation of the absolute value of the coefficients of all replications.

#### Epigenetic clock construction using ENR in the testing set

The CpG features were selected based on Pearson’s correlation coefficient between predicted DNAm age and chronological age in the testing set of MESA. Based on the CpG importance ranking, CpGs were added one at a time, and the correlation coefficients between predicted DNAm age and chronological age were calculated. We selected the CpG set with the correlation at the inflection point of the performance curve.

#### Evaluation of epigenetic clock performance in the validation dataset

The performance of the CpG features selected from MESA was evaluated in three independent validation cohorts (YSCCS, VACS, and WIHS) using the four measures, EAA, MDEAA, AMAR, and MDAMAR (see Statistics section).

### Statistics

#### Epigenetic clocks and assessments

Pearson correlation coefficients (r) between the DNAm age and chronological age were estimated in each cohort. EAA was defined as the residuals of regressing DNAm age on chronological age [5, 50]. We calculated the mean difference of EAA between heavy users and non-heavy users, between HIV+ and HIV-, and denoted it as MDEAA. AMAR was defined as the ratio of DNAm age to chronological age [6], AMAR>1 represents DNAm age acceleration, and AMAR<1 represents DNAm age deceleration. We also calculated the mean difference of AMAR between heavy and non-heavy users, between HIV+ and HIV-, and denoted as MDAMAR.

#### Benchmarking the MonoDNAmAge clock using HIV infection

We evaluated whether the MonoDNAmAge was able to predict changes in DNAm age for HIV infection in two cohorts (VACS and WIHS) using linear regression, EAA or AMAR as the dependent and HIV infection as the independent variable. We adjusted potential confounding factors including self-reported ethnicity, sex, tobacco use, body mass index (BMI), and alcohol-related phenotype in the model.

#### Association between alcohol consumption and DNAm-based biological age

Instead of just assuming a linear relationship between alcohol consumption and DNAm age, we first compared the linear model (reduced model) with the quadratic model (full model) for modeling the relationship between DNAm age and alcohol consumption (AUDIT in YSCCS; ln(PEth) in VACS; and NDRNKWK in WIHS) using ANOVA. The null hypothesis is that the reduced model is as good as the full model. We applied quadratic model when the quadratic model was significantly better than the linear model, otherwise, we applied the linear model.

In the quadratic regression model, DNAmAge=*β*_2_ *x*^2^ + *β*_1_*x* + *β*_0_, where *β*_2_ ≠ 0 and *x* represents alcohol consumption. The t-test was used to conduct hypothesis tests on the regression coefficients obtained in both quadratic regression and simple linear regression. That is, for the quadratic regression model, we tested *H*_0_: *β*_*j*_ = 0 vs. *β*_*j*_ ≠ 0 using t-test 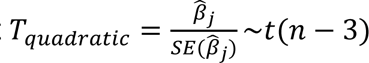, where *n* is the sample size. We also tested the regression model *H*_0_: the regression model was not significant vs. *H*_*α*_: the regression model was significant by performing F test, 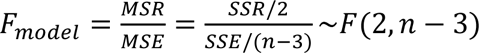 based on ANOVA test, where *MSR* was the regression mean square and *MSE* was the error mean square, 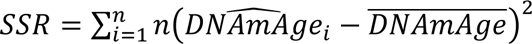 was the corrected sum of squares for regression model and 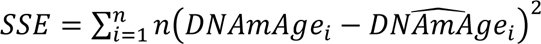 was the sum of squares for error.

We also examined EAA and AMAR between different groups of alcohol consumption in each cohort, adjusted potential confounding factors including self-reported ethnicity, sex, tobacco use, and BMI.

### Correlation between DNAm age and cell types in different alcohol consumption groups

To address potential cell type confounding effects, Pearson correlation coefficients (r) between DNAm-based estimates of biological age and six cell type proportions were estimated in each cohort.

## Acknowledgments

The authors appreciate the support of the Veterans Aging Study Cohort Biomarker Core (VACSBC), Yale Stress Center (YSC), Women’s Interagency HIV Study (WIHS), now the MACS/WIHS Combined Cohort Study (MWCCS), and the Yale Center of Genomic Analysis. Data in this manuscript were collected by the YSC, VACSBC, and WIHS/MWCCS. The contents of this publication are solely the responsibility of the authors and do not represent the official views of the National Institutes of Health (NIH).

## Conflict of Interest

The authors declare no conflict of interest.

## Funding

The project was supported by the National Institute on Drug Abuse [R03 DA039745 (Xu), R01 DA038632 (Xu), R01 DA047063 (Xu and Aouizerat), R01 DA047820(Xu and Aouizerat)], R01 DA013892 (Sinha), PL1 DA024859 (Sinha).

COMpAAAS/Veterans Aging Cohort Study, a CHAART Cooperative Agreement, supported by the National Institutes of Health: National Institute on Alcohol Abuse and Alcoholism (U24-AA020794, U01-AA020790, U01-AA020795, U01-AA020799; U10-AA013566-completed) and in kind by the US Department of Veterans Affairs. In addition to grant support from NIAAA, we gratefully acknowledge the scientific contributions of Dr. Kendall Bryant, our Scientific Collaborator. Additional grant support from National Institute on Drug Abuse R01-DA035616.

MWCCS (Principal Investigators): Atlanta CRS (Ighovwerha Ofotokun, Anandi Sheth, and Gina Wingood), U01-HL146241; Bronx CRS (Kathryn Anastos and Anjali Sharma), U01-HL146204; Brooklyn CRS (Deborah Gustafson and Tracey Wilson), U01-HL146202; Data Analysis and Coordination Center (Gypsyamber D’Souza, Stephen Gange and Elizabeth Golub), U01-HL146193; Chicago-Cook County CRS (Mardge Cohen and Audrey French), U01-HL146245; Northern California CRS (Bradley Aouizerat, Jennifer Price, and Phyllis Tien), U01-HL146242; Metropolitan Washington CRS (Seble Kassaye and Daniel Merenstein), U01-HL146205; Miami CRS (Maria Alcaide, Margaret Fischl, and Deborah Jones), U01-HL146203; UAB-MS CRS (Mirjam-Colette Kempf, Jodie Dionne-Odom, and Deborah Konkle-Parker), U01-HL146192; UNC CRS (Adaora Adimora), U01-HL146194. The MWCCS is funded primarily by the National Heart, Lung, and Blood Institute (NHLBI), with additional co-funding from the Eunice Kennedy Shriver National Institute Of Child Health & Human Development (NICHD), National Institute On Aging (NIA), National Institute Of Dental & Craniofacial Research (NIDCR), National Institute Of Allergy And Infectious Diseases (NIAID), National Institute Of Neurological Disorders And Stroke (NINDS), National Institute Of Mental Health (NIMH), National Institute On Drug Abuse (NIDA), National Institute Of Nursing Research (NINR), National Cancer Institute (NCI), National Institute on Alcohol Abuse and Alcoholism (NIAAA), National Institute on Deafness and Other Communication Disorders (NIDCD), National Institute of Diabetes and Digestive and Kidney Diseases (NIDDK), National Institute on Minority Health and Health Disparities (NIMHD), and in coordination and alignment with the research priorities of the National Institutes of Health, Office of AIDS Research (OAR). MWCCS data collection is also supported by UL1-TR000004 (UCSF CTSA), P30-AI-050409 (Atlanta CFAR), P30-AI-050410 (UNC CFAR), and P30-AI-027767 (UAB CFAR).

## Availability of Data and Materials

Demographic variables, clinical variables and methylation status for the VACS samples are submitted to the GEO dataset (GSE117861) and are available to the public. All codes for analysis are also available upon a request to the corresponding author.

## Authors’ Contributions

XL was responsible for bioinformatics data processing and statistical analysis. ACJ provided DNA samples and clinical data and contributed to the interpretation of findings and manuscript preparation. BA contributed to the design of the study. RS and BA provided DNA methylation and phenotype data, contributed to the interpretation of findings, and manuscript preparation. MC contributed to the manuscript preparation. KX was responsible for the study design, study protocol, sample preparation, data analysis, interpretation of findings, and manuscript preparation. All authors read and approved the final manuscript.

## Notes

### Competing Interest Statement

The authors have declared no competing interest.

