## Supplementary Information for "A New Monocyte Epigenetic Clock Reveals Effects of Alcohol Consumption on Epigenetic Aging in Three Independent Cohorts"

***All corresponding to**

Ke Xu, MD, PhD, Associate Professor of Psychiatry

Department of Psychiatry, Yale School of Medicine

VA Connecticut Healthcare System

950 Campbell Avenue

West Haven, CT 06516

**Study cohorts and phenotype assessment**

***Multi-Ethnic Study of Atherosclerosis (MESA) (N=1,202)*.** The MESA Epigenomics and Transcriptomics Study, funded by a National Heart, Lung, and Blood Institute, was established to investigate potential gene expression regulating methylation sites in human monocytes from a large study population [1]. The DNA samples in MESA (GSE56046) were from CD14+ monocyte samples, collected from 1,202 individuals with ages ranging from 44 to 83 years. The MESA cohort DNA methylation data was used solely to construct a monocyte-derived DNA methylation age clock using a machine learning approach.

***Yale Stress Center Cohort Study (YSCCS) (N=502)***. Five hundred and two healthy community volunteers were recruited from the community in the Greater New Haven area, Connecticut [2]. The participants who met the Diagnostic and Statistical Manual of Mental Disorders, 4th Edition (DSM-IVTR) (American Psychiatric Association, 1994) criteria for substance dependence on any drug other than nicotine were excluded to reduce confounding effects. Subjects with a diagnosis of medical illness or taking any medications for medical or psychiatric conditions were excluded. The cohort included men (44%) and women (56%) with ages ranging from 18 to 50 years. The participants included African Americans (AAs, 19.1%), European Americans (EAs, 72.2%), and other (8.7%). The cohort served as a community-based sample to examine alcohol consumption on epigenetic age among healthy participants. The 10-item Alcohol Use Disorders Identification Test (AUDIT) scale was used to assess alcohol use in YSCCS. Heavy alcohol drinking (HAD) (N=148) was defined as AUDIT≥8 for men and AUDIT≥7 for women based on previous studies while moderate alcohol drinking (MAD) (N=354) was defined as AUDIT<8 for men and AUDIT <7 for women [3]. The average AUDIT score was 5.69 among all participants in the cohort.

***Veterans Aging Cohort Study (VACS) (N=1,259).*** The VACS, funded primarily by the National Institute on Alcoholism and Alcohol Abuse, is an observational cohort study aiming to understand the role of comorbid medical and psychiatric disease in HIV infection in the United States [4]. Data from a subset of the cohort were collected through written consent, telephone interview, and access to the Veteran Affairs medical record system. Whole blood samples were collected within a similar window of time in conjunction with phenotypic data. The data available for analysis were generated from samples contributed by both HIV-positive (HIV+, N=1,151) and HIV-negative (HIV-, N=104) participants. All participants were AA men whose ages ranged from 25 to 79 years. The cohort served as a clinical-based sample to examine HIV infection and alcohol consumption on epigenetic age. Phosphatidylethanol (PEth) is a valid and clinically efficient biomarker for characterizing the recent alcohol use and differentiating light-moderate from heavy alcohol use and to corroborate self-report of alcohol use [5]. The levels of PEth measured in dried blood spots were positively correlated with AUDIT scores [6, 7]. The average PEth level in the VACS cohort was 41.67. HAD was defined as PEth≥20 (N=299, 29%) and MAD (N=738, 71%) was defined as PEth<20 in this cohort according to the previous studies [8].

***Women's Interagency HIV Study (WIHS) (N=481).*** The WIHS, established in 1993, is the longest-running multicenter, observational study of women with and at risk for HIV infection representative of the epidemic in the United States [9, 10]. The data available for analysis were generated from samples contributed by both HIV+ (N=272) and HIV

(N=209) participants, whose ages ranged from 18 to 78 years. The participants included AAs (49.1%), EAs (22.8%), and Hispanic Americans (28.1%). The cohort served as a clinical-based sample to examine HIV infection and alcohol consumption on epigenetic age. The self-reported number of drinks per week (NDRNKWK) was assessed for each participant at the same visit when blood samples were collected. The cohort predominantly reported light alcohol use with an average NDRNKWK of 0.7078. Light alcohol drinking (LAD) was defined as $0<\text{NDRNKWK}\leq 7$ (N=196, 43%), and non-alcohol drinking (NAD) was defined as NDRNKWK=0 (N=255, 57%) in this cohort.

Demographic and clinical variables for participants designated as HAD versus MAD, LAD versus NAD for each cohort were presented in **Table 1** in the article. Of note, low-moderate alcohol consumption was associated with less tobacco use, cannabis use, and less frequent alcohol intake compared to high alcohol consumption.

**Table S1. The set of CpGs selected by elastic net regularization for predicting MonoDNAmAge**

| Probe | | CHR | Position | Gene | Group | p | FDR | Coefficient |
|---|---|---|---|---|---|---|---|---|
| 1 | cg11220950 | 16 | 2042693 | *SYNGR3* | Body | 9.64E-39 | 3.77E-35 | 28.27 |
| 2 | cg26685941 | 13 | 95952902 | *ABCC4* | Body | 8.51E-37 | 2.61E-33 | -23.26 |
| 3 | cg21186299 | 7 | 100808810 | *VGF* | 5UTR | 7.71E-27 | 6.63E-24 | 22.36 |
| 4 | cg22158769 | 2 | 39187539 | *LOC375196; LOC100271715* | TSS200 | 2.27E-74 | 3.86E-69 | 27.44 |
| 5 | cg12623930 | 3 | 52008802 | *ABHD14B; ABHD14A* | TSS1500 | 4.44E-38 | 1.57E-34 | -20.59 |
| 6 | cg08295410 | 5 | 156990663 | *ADAM19* | Body | 1.37E-28 | 1.49E-25 | -26.83 |
| 7 | cg06493994 | 6 | 25652602 | *SCGN* | 5UTR | 1.11E-58 | 4.19E-54 | 20.09 |
| 8 | cg09499629 | 7 | 130419136 | *KLF14* | TSS1500 | 7.38E-40 | 3.22E-36 | 16.45 |
| 9 | cg22285878 | 7 | 130419173 | *KLF14* | TSS1500 | 2.12E-46 | 2.06E-42 | 19.31 |
| 10 | cg25157472 | 3 | 12996362 | *IQSEC1* | Body | 5.49E-30 | 7.15E-27 | 22.56 |
| 11 | cg09726279 | 17 | 4457835 | *MYBBP1A* | Body | 2.55E-20 | 9.07E-18 | -21.67 |
| 12 | cg00281467 | 22 | 23522460 | *BCR* | TSS200 | 5.53E-25 | 3.71E-22 | -17.13 |
| 13 | cg09893871 | 6 | 166720997 | *PRR18* | 1stExon | 8.11E-34 | 1.58E-30 | 12.60 |
| 14 | cg07171111 | 4 | 10462903 | NA | NA | 6.18E-26 | 4.76E-23 | 17.04 |
| 15 | cg04434593 | 11 | 67139546 | *LOC100130987; CLCF1* | 5UTR | 1.39E-22 | 6.72E-20 | -18.03 |
| 16 | cg13483882 | 3 | 128212476 | *GATA2* | TSS1500 | 5.89E-23 | 3.01E-20 | -14.65 |
| 17 | cg10778288 | 12 | 113917994 | NA | NA | 8.20E-25 | 5.35E-22 | 15.05 |
| 18 | cg22557662 | 19 | 38747374 | *PPP1R14A* | TSS1500 | 6.98E-36 | 1.85E-32 | 16.48 |
| 19 | cg00745389 | 7 | 32467435 | NA | NA | 3.10E-67 | 3.51E-62 | 13.42 |
| 20 | cg19401340 | 17 | 56833197 | *PPM1E* | TSS200 | 1.21E-26 | 1.03E-23 | 15.31 |
| 21 | cg21166964 | 5 | 72529816 | NA | NA | 3.09E-22 | 1.42E-19 | 14.92 |
| 22 | cg05713859 | 5 | 134240166 | *PCBD2* | TSS1500 | 3.84E-24 | 2.29E-21 | -18.84 |
| 23 | cg22565251 | 19 | 45461093 | *CLPTM1* | Body | 1.64E-20 | 6.01E-18 | -20.93 |
| 24 | cg07098391 | 4 | 55991418 | *KDR* | 1stExon | 4.73E-23 | 2.44E-20 | 16.79 |
| 25 | cg04501188 | 1 | 47904171 | *FOXD2* | 1stExon | 3.40E-44 | 2.31E-40 | 13.15 |
| 26 | cg04940570 | 11 | 12696758 | *TEAD1* | 5UTR | 1.98E-50 | 2.80E-46 | 14.00 |
| 27 | cg21383487 | 10 | 22624094 | NA | NA | 2.39E-21 | 9.78E-19 | 11.99 |
| 28 | cg25726357 | 6 | 163147733 | *PACRG; PARK2* | TSS1500 | 2.14E-21 | 8.83E-19 | -19.47 |
| 29 | cg03930964 | 22 | 23522374 | *BCR* | TSS200 | 6.17E-32 | 9.89E-29 | -13.07 |
| 30 | cg00034076 | 8 | 85096037 | *RALYL* | TSS1500 | 2.64E-27 | 2.43E-24 | 12.78 |
| 31 | cg01907194 | 14 | 104003370 | *TRMT61A* | 3UTR | 1.41E-19 | 4.60E-17 | 12.46 |
| 32 | cg06416491 | 12 | 39837660 | *KIF21A* | TSS1500 | 3.60E-56 | 8.74E-52 | -12.15 |
| 33 | cg09295081 | 2 | 11810183 | *NTSR2* | 1stExon | 9.12E-46 | 7.62E-42 | 12.43 |
| 34 | cg07254032 | 5 | 45695839 | *HCN1* | 1stExon | 2.76E-26 | 2.23E-23 | 15.54 |
| 35 | cg21283081 | 3 | 128152179 | NA | NA | 1.95E-21 | 8.10E-19 | -10.23 |
| 36 | cg11299854 | 5 | 132083184 | *CCNI2* | 5UTR | 4.55E-21 | 1.78E-18 | -17.59 |
| 37 | cg07705835 | 3 | 9959128 | *IL17RC* | 1stExon | 2.06E-27 | 1.95E-24 | 13.70 |
| 38 | cg06943835 | 11 | 64662577 | *ATG2A* | Body | 2.17E-36 | 6.25E-33 | -13.11 |
| 39 | cg00885918 | 20 | 30406997 | *MYLK2* | TSS200 | 4.79E-26 | 3.76E-23 | 16.73 |
| 40 | cg12422450 | 14 | 93389891 | *CHGA* | Body | 2.18E-27 | 2.05E-24 | 10.27 |
| 41 | cg13203811 | 12 | 58136245 | *AGAP2* | TSS1500 | 2.50E-34 | 5.28E-31 | 15.41 |

| | | | | | | | | |
|---|---|---|---|---|---|---|---|---|
| 42 | cg20346726 | 11 | 101918403 | *C11orf70* | 5UTR | 2.77E-21 | 1.12E-18 | -18.05 |
| 43 | cg14912644 | 2 | 157176601 | NA | NA | 1.87E-21 | 7.79E-19 | 9.17 |
| 44 | cg12189835 | 11 | 61335071 | *SYT7* | Body | 5.14E-28 | 5.15E-25 | 11.71 |
| 45 | cg18363008 | 17 | 4675114 | *TM4SF5* | TSS200 | 1.09E-40 | 5.15E-37 | -14.78 |
| 46 | cg23346544 | 8 | 80731144 | NA | NA | 4.54E-20 | 1.57E-17 | -12.25 |
| 47 | cg12446246 | 1 | 208418552 | *PLXNA2* | TSS1500 | 3.94E-27 | 3.52E-24 | -11.21 |
| 48 | cg16081281 | 8 | 23559839 | NA | NA | 4.93E-21 | 1.92E-18 | 11.67 |
| 49 | cg12920180 | 14 | 31344444 | *COCH* | Body | 1.80E-29 | 2.19E-26 | 11.17 |
| 50 | cg17396222 | 7 | 134228596 | NA | NA | 5.60E-28 | 5.58E-25 | -13.61 |
| 51 | cg11159604 | 1 | 91316223 | NA | NA | 1.16E-25 | 8.59E-23 | -13.29 |
| 52 | cg13053396 | 12 | 7168545 | *C1S* | 5UTR | 3.68E-20 | 1.29E-17 | -13.28 |
| 53 | cg26325867 | 6 | 41398693 | NA | NA | 7.61E-41 | 3.75E-37 | -12.90 |
| 54 | cg26062560 | 3 | 194979832 | *C3orf21* | Body | 7.53E-20 | 2.53E-17 | 10.97 |
| 55 | cg10947146 | 8 | 11058710 | *XKR6* | 1stExon | 3.49E-36 | 9.64E-33 | 8.34 |
| 56 | cg04673446 | 22 | 39879951 | *MGAT3* | 5UTR | 4.79E-21 | 1.87E-18 | -12.22 |
| 57 | cg16193278 | 13 | 100008450 | *UBAC2; MIR623* | Body | 7.36E-45 | 5.44E-41 | -13.32 |
| 58 | cg10576725 | 14 | 23306974 | *MMP14* | Body | 2.06E-28 | 2.17E-25 | -13.07 |
| 59 | cg03605420 | 5 | 74162809 | *FAM169A* | TSS200 | 1.00E-40 | 4.79E-37 | -8.28 |
| 60 | cg07571951 | 7 | 75931608 | *HSPB1* | TSS1500 | 1.48E-25 | 1.08E-22 | -13.21 |
| 61 | cg26158023 | 3 | 42881414 | *CCBP2* | 5UTR | 1.27E-36 | 3.72E-33 | -13.46 |
| 62 | cg03335216 | 1 | 8086776 | *ERRFI1* | TSS1500 | 5.80E-31 | 8.35E-28 | 8.35 |
| 63 | cg02383785 | 7 | 127808848 | NA | NA | 1.22E-34 | 2.71E-31 | 7.90 |
| 64 | cg01981760 | 16 | 53737576 | *FTO; RPGRIP1L* | TSS1500 | 1.38E-23 | 7.64E-21 | -10.53 |
| 65 | cg02018902 | 15 | 79576149 | *ANKRD34C* | 5UTR | 5.46E-21 | 2.12E-18 | 8.88 |
| 66 | cg06815715 | 15 | 49148520 | *SHC4* | Body | 4.38E-38 | 1.57E-34 | 14.13 |
| 67 | cg26864395 | 1 | 25259106 | *RUNX3* | Body | 1.95E-34 | 4.22E-31 | 17.50 |
| 68 | cg06193004 | 20 | 43729874 | *KCNS1* | TSS200 | 2.39E-34 | 5.08E-31 | -9.85 |
| 69 | cg01787559 | 4 | 3450770 | *HGFAC* | Body | 9.13E-25 | 5.91E-22 | 12.98 |
| 70 | cg06872770 | 1 | 3688128 | *LOC388588; CCDC27* | TSS1500 | 2.02E-21 | 8.37E-19 | -12.87 |
| 71 | cg03650713 | 1 | 16501734 | NA | NA | 6.58E-21 | 2.53E-18 | 12.38 |
| 72 | cg07639287 | 22 | 17850911 | NA | NA | 1.40E-22 | 6.75E-20 | 9.32 |
| 73 | cg00116092 | 1 | 217313044 | NA | NA | 7.12E-43 | 4.48E-39 | 3.55 |
| 74 | cg03211864 | 10 | 124060833 | *BTBD16* | Body | 4.72E-20 | 1.63E-17 | -11.41 |
| 75 | cg24632582 | 15 | 41233701 | NA | NA | 7.75E-23 | 3.87E-20 | -13.23 |
| 76 | cg16881676 | 12 | 49659619 | *TUBA1C* | Body | 1.44E-22 | 6.93E-20 | -13.92 |
| 77 | cg08739433 | 1 | 113051925 | *WNT2B* | 1stExon | 4.91E-41 | 2.49E-37 | 5.98 |
| 78 | cg01807026 | 9 | 123475367 | *MEGF9* | Body | 2.20E-30 | 3.00E-27 | -7.11 |
| 79 | cg24013954 | 7 | 140394679 | *ADCK2* | 3UTR | 1.47E-22 | 7.05E-20 | -7.97 |
| 80 | cg27376872 | 8 | 899171 | NA | NA | 1.30E-22 | 6.30E-20 | 16.06 |
| 81 | cg05056653 | 17 | 81023856 | NA | NA | 9.30E-20 | 3.09E-17 | 16.44 |
| 82 | cg24517738 | 1 | 41129755 | *RIMS3* | 5UTR | 3.53E-22 | 1.61E-19 | 12.07 |
| 83 | cg00318320 | 2 | 242498098 | *BOK* | TSS200 | 9.98E-35 | 2.25E-31 | 11.12 |
| 84 | cg21922223 | 17 | 75539118 | NA | NA | 5.71E-35 | 1.34E-31 | -11.78 |
| 85 | cg00982799 | 10 | 129868087 | *PTPRE* | Body | 2.49E-26 | 2.02E-23 | -11.22 |
| 86 | cg21801378 | 15 | 72612125 | *BRUNOL6* | 1stExon | 4.53E-43 | 2.96E-39 | 5.75 |

| | | | | | | | | |
|---|---|---|---|---|---|---|---|---|
| 87 | cg01797043 | 16 | 2004686 | *RPL3L* | TSS200 | 2.92E-56 | 7.63E-52 | -12.32 |
| 88 | cg20445053 | 2 | 169440359 | *LASS6* | Body | 1.99E-36 | 5.78E-33 | -11.12 |
| 89 | cg04967200 | 10 | 134115376 | *STK32C* | Body | 1.37E-23 | 7.61E-21 | 9.84 |
| 90 | cg14999189 | 17 | 2908369 | *RAP1GAP2* | Body | 1.36E-27 | 1.32E-24 | 7.14 |
| 91 | cg06640584 | 2 | 227700558 | *RHBDD1* | TSS200 | 2.22E-27 | 2.08E-24 | 5.33 |
| 92 | cg02446869 | 6 | 31654389 | NA | NA | 6.95E-30 | 8.88E-27 | -9.43 |
| 93 | cg16998353 | 11 | 6495553 | *TRIM3* | TSS1500 | 4.78E-35 | 1.14E-31 | -10.70 |
| 94 | cg02867102 | 17 | 62398693 | NA | NA | 8.14E-40 | 3.50E-36 | -5.49 |
| 95 | cg27213509 | 2 | 176947228 | *EVX2* | Body | 2.10E-24 | 1.30E-21 | 8.52 |
| 96 | cg21523251 | 4 | 8582110 | *GPR78* | TSS200 | 8.66E-26 | 6.53E-23 | 6.87 |
| 97 | cg00344380 | 4 | 42155519 | *BEND4* | TSS1500 | 4.02E-32 | 6.57E-29 | -14.91 |
| 98 | cg16006701 | 6 | 2751402 | *MYLK4* | TSS1500 | 3.06E-25 | 2.14E-22 | -10.72 |
| 99 | cg10151162 | 16 | 333104 | *PDIA2* | TSS200 | 6.00E-30 | 7.73E-27 | -7.68 |
| 100 | cg21505886 | 4 | 1724428 | *TMEM129; TACC3* | TSS1500 | 1.02E-21 | 4.38E-19 | 11.61 |
| 101 | cg13576991 | 1 | 223257466 | NA | NA | 8.65E-20 | 2.89E-17 | -3.84 |
| 102 | cg16744250 | 20 | 62209504 | NA | NA | 2.52E-22 | 1.18E-19 | -7.96 |
| 103 | cg05308819 | 1 | 155959156 | NA | NA | 3.18E-42 | 1.83E-38 | -5.25 |
| 104 | cg08822715 | 12 | 114404604 | *RBM19* | TSS1500 | 5.94E-20 | 2.03E-17 | -7.28 |
| 105 | cg24155190 | 1 | 201476619 | *CSRP1* | TSS1500 | 4.24E-21 | 1.67E-18 | -10.05 |
| 106 | cg09790829 | 6 | 91007365 | *BACH2* | TSS1500 | 3.83E-22 | 1.73E-19 | 7.93 |
| 107 | cg24170090 | 17 | 16945301 | *MPRIP* | TSS1500 | 7.71E-28 | 7.62E-25 | -11.76 |
| 108 | cg01230796 | 6 | 25652698 | *SCGN* | 1stExon | 1.28E-22 | 6.22E-20 | 8.01 |
| 109 | cg13649056 | 9 | 136474626 | NA | NA | 5.67E-82 | 1.93E-76 | 8.36 |
| 110 | cg23495995 | 11 | 65311181 | *LTBP3* | Body | 8.91E-25 | 5.79E-22 | 13.15 |
| 111 | cg03443986 | 2 | 65100572 | NA | NA | 9.74E-47 | 1.00E-42 | -7.28 |
| 112 | cg24853724 | 7 | 28997403 | *TRIL* | 1stExon | 1.18E-47 | 1.34E-43 | 6.72 |
| 113 | cg26842024 | 19 | 16436122 | *KLF2* | Body | 1.74E-24 | 1.09E-21 | 7.24 |
| 114 | cg03664992 | 1 | 39957393 | *BMP8A* | 5UTR | 9.39E-34 | 1.82E-30 | 6.86 |
| 115 | cg22786465 | 6 | 31649502 | *LY6G5C* | TSS1500 | 6.68E-20 | 2.26E-17 | 11.52 |
| 116 | cg16567172 | 7 | 75931606 | *HSPB1* | TSS1500 | 3.20E-26 | 2.56E-23 | -9.68 |
| 117 | cg14755852 | 8 | 23386014 | *SLC25A37* | TSS1500 | 2.84E-23 | 1.50E-20 | -8.52 |
| 118 | cg23994468 | 19 | 56147306 | NA | NA | 1.60E-22 | 7.64E-20 | -11.56 |
| 119 | cg22833065 | 17 | 38095691 | NA | NA | 4.99E-20 | 1.72E-17 | -7.08 |
| 120 | cg00484358 | 1 | 110610995 | *ALX3* | Body | 1.47E-23 | 8.10E-21 | 7.06 |
| 121 | cg04084157 | 7 | 100809049 | *VGF* | TSS200 | 1.30E-33 | 2.47E-30 | 7.84 |
| 122 | cg13347071 | 2 | 210636748 | *UNC80* | 5UTR | 8.88E-35 | 2.01E-31 | 8.52 |
| 123 | cg24351658 | 1 | 42126724 | *HIVEP3* | 5UTR | 4.80E-28 | 4.83E-25 | 10.43 |
| 124 | cg11864592 | 15 | 85833204 | NA | NA | 4.42E-20 | 1.54E-17 | 11.63 |
| 125 | cg06482428 | 4 | 21950173 | *KCNIP4* | 5UTR | 3.05E-26 | 2.46E-23 | -6.90 |
| 126 | cg12774845 | 14 | 74486312 | *C14orf45; ENTPD5* | TSS1500 | 1.65E-24 | 1.03E-21 | 10.54 |
| 127 | cg07508304 | 16 | 85208316 | NA | NA | 1.86E-20 | 6.74E-18 | 17.17 |
| 128 | cg00296038 | 3 | 11624533 | *VGLL4* | TSS1500 | 1.07E-19 | 3.53E-17 | 10.42 |
| 129 | cg14538537 | 12 | 6939048 | *LEPREL2* | Body | 4.91E-20 | 1.69E-17 | 4.59 |
| 130 | cg08761208 | 15 | 65693289 | *IGDCC4* | Body | 6.22E-66 | 4.23E-61 | -7.87 |
| 131 | cg11215976 | 1 | 236849942 | *ACTN2* | 5UTR | 2.02E-23 | 1.09E-20 | 8.15 |
| 132 | cg23917779 | 1 | 36807505 | *STK40* | Body | 3.39E-25 | 2.35E-22 | -7.94 |

| | | | | | | | | |
|---|---|---|---|---|---|---|---|---|
| 133 | cg25826226 | 15 | 41953061 | *MGA* | 5UTR | 3.13E-33 | 5.78E-30 | 5.88 |
| 134 | cg18299578 | 14 | 29235928 | *FOXG1* | TSS1500 | 1.27E-25 | 9.32E-23 | 9.93 |
| 135 | cg00081324 | 17 | 48201233 | *SAMD14* | Body | 9.08E-27 | 7.74E-24 | 14.34 |
| 136 | cg17133388 | 3 | 122102727 | *FAM162A; CCDC58* | TSS1500 | 6.12E-25 | 4.05E-22 | 8.54 |
| 137 | cg27075187 | 11 | 12696059 | *TEAD1* | 5UTR | 4.21E-21 | 1.66E-18 | 6.62 |
| 138 | cg15821095 | 3 | 51428079 | *RBM15B* | TSS1500 | 2.84E-21 | 1.14E-18 | 13.55 |
| 139 | cg07092212 | 11 | 46382544 | *DGKZ* | TSS1500 | 6.86E-36 | 1.84E-32 | -4.14 |
| 140 | cg08124461 | 15 | 57818971 | *CGNL1* | Body | 5.05E-22 | 2.24E-19 | -8.04 |
| 141 | cg24812167 | 9 | 139293358 | *SNAPC4* | TSS1500 | 2.54E-21 | 1.03E-18 | 10.08 |
| 142 | cg00109344 | 9 | 116344532 | *RGS3* | 5UTR | 5.23E-24 | 3.08E-21 | 10.05 |
| 143 | cg06685111 | 6 | 30295466 | *HCG18; TRIM39* | TSS1500 | 1.06E-36 | 3.19E-33 | -8.90 |
| 144 | cg25618916 | 1 | 10825554 | *CASZ1* | 5UTR | 1.14E-19 | 3.75E-17 | -6.62 |
| 145 | cg04596060 | 12 | 114404600 | *RBM19* | TSS1500 | 1.51E-19 | 4.90E-17 | -8.68 |
| 146 | cg16989340 | 1 | 1084147 | NA | NA | 7.21E-23 | 3.63E-20 | -8.28 |
| 147 | cg07718813 | 1 | 6640976 | *ZBTB48* | Body | 2.58E-22 | 1.20E-19 | 9.27 |
| 148 | cg27449864 | 7 | 73133982 | *STX1A* | 5UTR | 2.16E-24 | 1.33E-21 | 7.93 |
| 149 | cg16127845 | 7 | 1126423 | *GPER; C7orf50* | TSS1500 | 2.53E-25 | 1.81E-22 | 11.93 |
| 150 | cg19442470 | 8 | 27470225 | *CLU* | TSS1500 | 3.26E-38 | 1.18E-34 | 11.83 |
| 151 | cg10257332 | 3 | 47869137 | *DHX30* | Body | 2.65E-46 | 2.50E-42 | -3.02 |
| 152 | cg04673912 | 7 | 116593113 | *ST7OT4; ST7OT1; ST7* | TSS1500 | 3.63E-30 | 4.80E-27 | -6.04 |
| 153 | cg00653615 | 1 | 151029327 | *CDC42SE1* | 5UTR | 1.21E-23 | 6.76E-21 | 10.34 |
| 154 | cg01526854 | 6 | 41900498 | *BYSL* | 3UTR | 5.38E-31 | 7.78E-28 | -11.08 |
| 155 | cg18724565 | 7 | 5633068 | *FSCN1* | 1stExon | 2.50E-32 | 4.25E-29 | 7.11 |
| 156 | cg18292202 | 6 | 26746947 | NA | NA | 8.09E-20 | 2.71E-17 | -11.58 |
| 157 | cg11905007 | 3 | 62861071 | *CADPS* | TSS200 | 3.49E-20 | 1.23E-17 | 7.83 |
| 158 | cg10713715 | 11 | 63533656 | *C11orf95* | Body | 1.12E-32 | 1.99E-29 | -6.66 |
| 159 | cg21574064 | 19 | 8661935 | *ADAMTS10* | Body | 1.32E-20 | 4.89E-18 | -10.89 |
| 160 | cg14564815 | 5 | 139752588 | *SLC4A9* | 3UTR | 3.58E-28 | 3.70E-25 | -9.85 |
| 161 | cg00384539 | 8 | 70983567 | *PRDM14* | TSS200 | 5.48E-27 | 4.83E-24 | 9.05 |
| 162 | cg09547767 | 1 | 39957387 | *BMP8A* | 5UTR | 8.49E-24 | 4.80E-21 | 7.95 |
| 163 | cg02415029 | 1 | 110025642 | *ATXN7L2* | TSS1500 | 1.17E-28 | 1.30E-25 | -6.06 |
| 164 | cg26231732 | 10 | 100069542 | NA | NA | 1.62E-24 | 1.02E-21 | -5.32 |
| 165 | cg25307771 | 13 | 91067121 | NA | NA | 6.11E-25 | 4.05E-22 | -2.90 |
| 166 | cg25930644 | 17 | 8531915 | *MYH10* | 5UTR | 3.82E-32 | 6.27E-29 | 3.83 |
| 167 | cg10117719 | 1 | 36181173 | *C1orf216* | 3UTR | 9.20E-29 | 1.03E-25 | -10.88 |
| 168 | cg15428620 | 10 | 102792835 | *SFXN3* | Body | 1.71E-25 | 1.24E-22 | -4.24 |
| 169 | cg15370180 | 10 | 23484326 | NA | NA | 7.34E-20 | 2.47E-17 | 7.53 |
| 170 | cg02933228 | 11 | 64611087 | *CDC42BPG* | Body | 2.08E-21 | 8.60E-19 | -5.73 |
| 171 | cg14518803 | 5 | 175662789 | NA | NA | 2.63E-20 | 9.31E-18 | 12.07 |
| 172 | cg27151122 | 12 | 50447407 | NA | NA | 3.22E-32 | 5.39E-29 | -5.43 |
| 173 | cg23460950 | 8 | 27472513 | *CLU* | TSS200 | 7.40E-30 | 9.35E-27 | -7.80 |
| 174 | cg25353142 | 20 | 43729869 | *KCNS1* | TSS200 | 2.61E-25 | 1.86E-22 | -2.68 |
| 175 | cg25533247 | 19 | 15530630 | *AKAP8L* | TSS1500 | 5.87E-42 | 3.22E-38 | -4.15 |
| 176 | cg11911550 | 5 | 39716771 | NA | NA | 3.51E-28 | 3.64E-25 | 13.65 |
| 177 | cg12552835 | 19 | 11032057 | *CARM1* | Body | 1.42E-21 | 6.02E-19 | 8.74 |

| | | | | | | | | |
|---|---|---|---|---|---|---|---|---|
| 178 | cg07558287 | 3 | 127347884 | *PODXL2* | TSS200 | 4.37E-28 | 4.45E-25 | 9.37 |
| 179 | cg20974724 | 7 | 121944450 | *FEZF1* | 1stExon | 8.09E-21 | 3.09E-18 | 6.80 |
| 180 | cg05331060 | 6 | 31459999 | NA | NA | 1.44E-33 | 2.72E-30 | -9.21 |
| 181 | cg10801607 | 2 | 44518930 | *SLC3A1* | Body | 1.19E-20 | 4.44E-18 | 6.54 |
| 182 | cg02807849 | 19 | 48908102 | *GRIN2D* | Body | 3.65E-27 | 3.29E-24 | -4.20 |
| 183 | cg01812045 | 2 | 220283739 | *DES* | 1stExon | 1.08E-25 | 8.03E-23 | 4.38 |
| 184 | cg06196801 | 12 | 121164025 | *ACADS* | Body | 1.09E-29 | 1.35E-26 | -4.59 |
| 185 | cg25570486 | 20 | 61621446 | NA | NA | 5.97E-23 | 3.04E-20 | -8.28 |
| 186 | cg17051207 | 12 | 49482508 | NA | NA | 2.18E-28 | 2.29E-25 | -12.22 |

MonoDNAmAge: DNA methylation monocyte age; CHR: chromosome; FDR: Benjamini-Hochberg false discovery rate adjusted P value; Coefficient: coefficient of linear regression analysis.

**Table S2. Association between EAA/AMAR and HIV**

| Measure | Cohort | Method | beta | 95% CI | p | MD |
|---|---|---|---|---|---|---|
| EAA | YSCCS | MonoDNAmAge | NA | NA | NA | NA |
| | | HorvathDNAmAge | NA | NA | NA | NA |
| | | HannumDNAmAge | NA | NA | NA | NA |
| | | PhenoDNAmAge | NA | NA | NA | NA |
| | | GrimDNAmAge | NA | NA | NA | NA |
| | VACS | MonoDNAmAge | 9.972 | (8.107, 11.838) | **1.17E-24** | 10.139 |
| | | HorvathDNAmAge | 2.931 | (1.763, 4.099) | **9.90E-07** | 2.527 |
| | | HannumDNAmAge | -3.267 | (-4.449, -2.085) | **7.29E-08** | -3.432 |
| | | PhenoDNAmAge | 1.972 | (0.459, 3.484) | **1.07E-02** | 2.365 |
| | | GrimDNAmAge | -4.391 | (-5.676, -3.106) | **3.26E-11** | -3.463 |
| | WIHS | MonoDNAmAge | 12.014 | (10.021, 14.007) | **2.07E-28** | 12.172 |
| | | HorvathDNAmAge | 3.580 | (2.678, 4.482) | **4.62E-14** | 3.603 |
| | | HannumDNAmAge | 2.886 | (1.731, 4.042) | **1.34E-06** | 3.038 |
| | | PhenoDNAmAge | 5.339 | (3.962, 6.716) | **1.61E-13** | 5.199 |
| | | GrimDNAmAge | 1.323 | (0.650, 1.996) | **1.34E-04** | 0.851 |
| AMAR | YSCCS | MonoDNAmAge | NA | NA | NA | NA |
| | | HorvathDNAmAge | NA | NA | NA | NA |
| | | HannumDNAmAge | NA | NA | NA | NA |
| | | PhenoDNAmAge | NA | NA | NA | NA |
| | | GrimDNAmAge | NA | NA | NA | NA |
| | VACS | MonoDNAmAge | 0.209 | (0.172, 0.245) | **1.00E-27** | 0.209 |
| | | HorvathDNAmAge | 0.064 | (0.039, 0.090) | **1.01E-06** | 0.064 |
| | | HannumDNAmAge | -0.055 | (-0.080, -0.030) | **2.09E-05** | -0.055 |
| | | PhenoDNAmAge | 0.058 | (0.027, 0.089) | **2.74E-04** | 0.058 |
| | | GrimDNAmAge | -0.047 | (-0.080, -0.014) | **4.91E-03** | -0.047 |
| | WIHS | MonoDNAmAge | 0.341 | (0.293, 0.389) | **6.51E-37** | 0.341 |
| | | HorvathDNAmAge | 0.079 | (0.055, 0.102) | **7.13E-11** | 0.079 |
| | | HannumDNAmAge | 0.052 | (0.021, 0.082) | **8.63E-04** | 0.052 |
| | | PhenoDNAmAge | 0.177 | (0.139, 0.214) | **9.01E-19** | 0.177 |
| | | GrimDNAmAge | -0.010 | (-0.031, 0.010) | 3.17E-01 | -0.010 |

EAA: Epigenetic Age Acceleration; AMAR: Apparent Methylation Age Rate.
YSCCS: Yale Stress Center Cohort Study; VACS: Veterans Aging Cohort Study; WIHS: Women's Interagency HIV Study.
95% CI: 95% confidence interval for coefficient beta.
The p-values in bold indicate the values less than 5.00E-02.

**Table S3. P-values of model comparison between the linear model and quadratic model using ANOVA**

| Method | ANOVA F-statistic p-value | | |
|---|---|---|---|
| | **YSCCS** | **VACS** | **WIHS** |
| MonoDNAmAge | **7.80E-08** | **3.46E-02** | 2.47E-01 |
| HorvathDNAmAge | **2.37E-07** | **3.91E-02** | 3.17E-01 |
| HannumDNAmAge | **2.41E-06** | **8.32E-04** | 4.38E-01 |
| PhenoDNAmAge | **6.27E-09** | **8.60E-04** | 2.72E-01 |
| GrimDNAmAge | **6.12E-08** | **2.21E-02** | 3.38E-01 |

The p-values in bold indicate the values less than 5.00E-02.

**(a)**

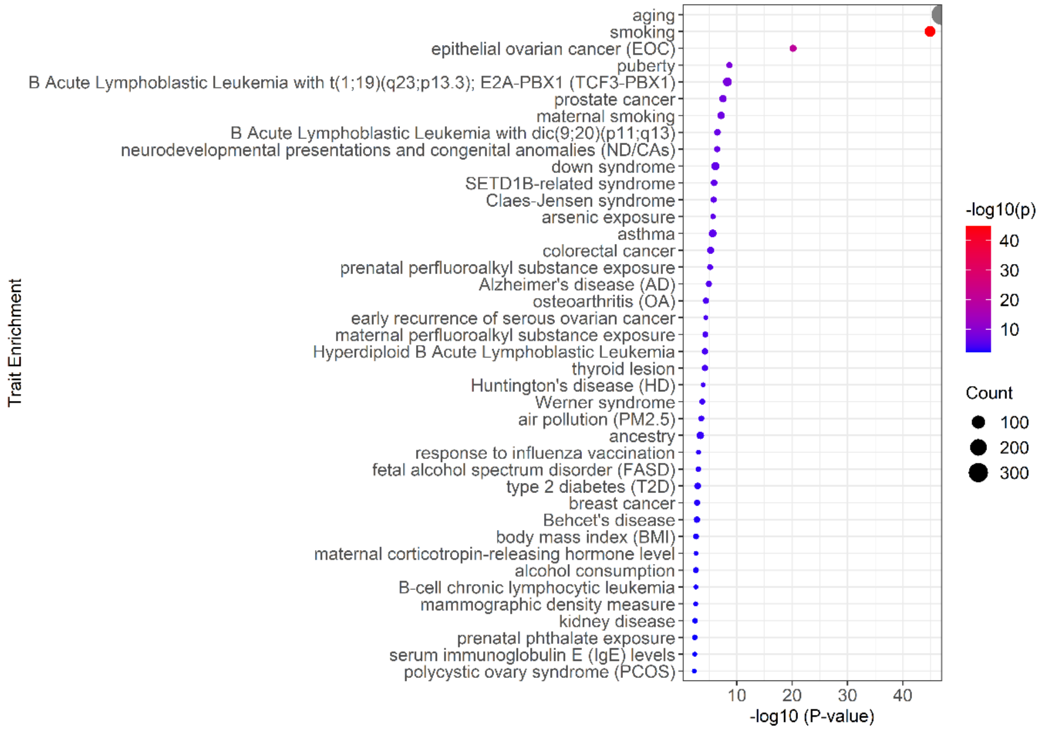

aging
smoking
epithelial ovarian cancer (EOC)
puberty
B Acute Lymphoblastic Leukemia with t(1;19)(q23;p13.3); E2A-PBX1 (TCF3-PBX1)
prostate cancer
maternal smoking
B Acute Lymphoblastic Leukemia with dic(9;20)(p11;q13)
neurodevelopmental presentations and congenital anomalies (ND/CAs)
down syndrome
SETD1B-related syndrome
Claes-Jensen syndrome
arsenic exposure
asthma
colorectal cancer
prenatal perfluoroalkyl substance exposure
Alzheimer's disease (AD)
osteoarthritis (OA)
early recurrence of serous ovarian cancer
maternal perfluoroalkyl substance exposure
Hyperdiploid B Acute Lymphoblastic Leukemia
thyroid lesion
Huntington's disease (HD)
Werner syndrome
air pollution (PM2.5)
ancestry
response to influenza vaccination
fetal alcohol spectrum disorder (FASD)
type 2 diabetes (T2D)
breast cancer
Behcet's disease
body mass index (BMI)
maternal corticotropin-releasing hormone level
alcohol consumption
B-cell chronic lymphocytic leukemia
mammographic density measure
kidney disease
prenatal phthalate exposure
serum immunoglobulin E (IgE) levels
polycystic ovary syndrome (PCOS)
Trait Enrichment
10 20 30 40
-log10 (P-value)
-log10(p)
40
30
20
10
Count
100
200
300

**(b)**

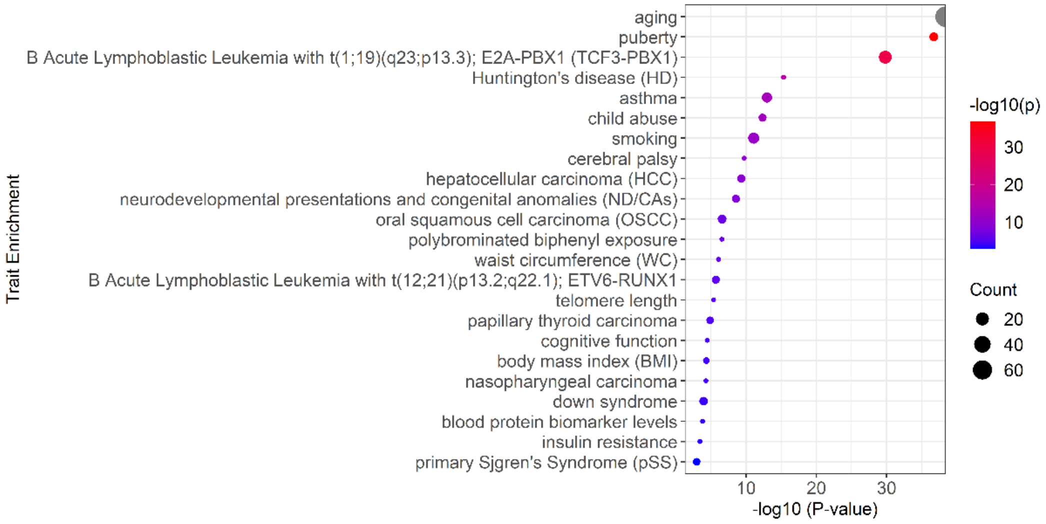

aging
puberty
B Acute Lymphoblastic Leukemia with t(1;19)(q23;p13.3); E2A-PBX1 (TCF3-PBX1)
Huntington's disease (HD)
asthma
child abuse
smoking
cerebral palsy
hepatocellular carcinoma (HCC)
neurodevelopmental presentations and congenital anomalies (ND/CAs)
oral squamous cell carcinoma (OSCC)
polybrominated biphenyl exposure
waist circumference (WC)
B Acute Lymphoblastic Leukemia with t(12;21)(p13.2;q22.1); ETV6-RUNX1
telomere length
papillary thyroid carcinoma
cognitive function
body mass index (BMI)
nasopharyngeal carcinoma
down syndrome
blood protein biomarker levels
insulin resistance
primary Sjgren's Syndrome (pSS)
Trait Enrichment
10 20 30
-log10 (P-value)
-log10(p)
30
20
10
Count
20
40
60

**(c)**

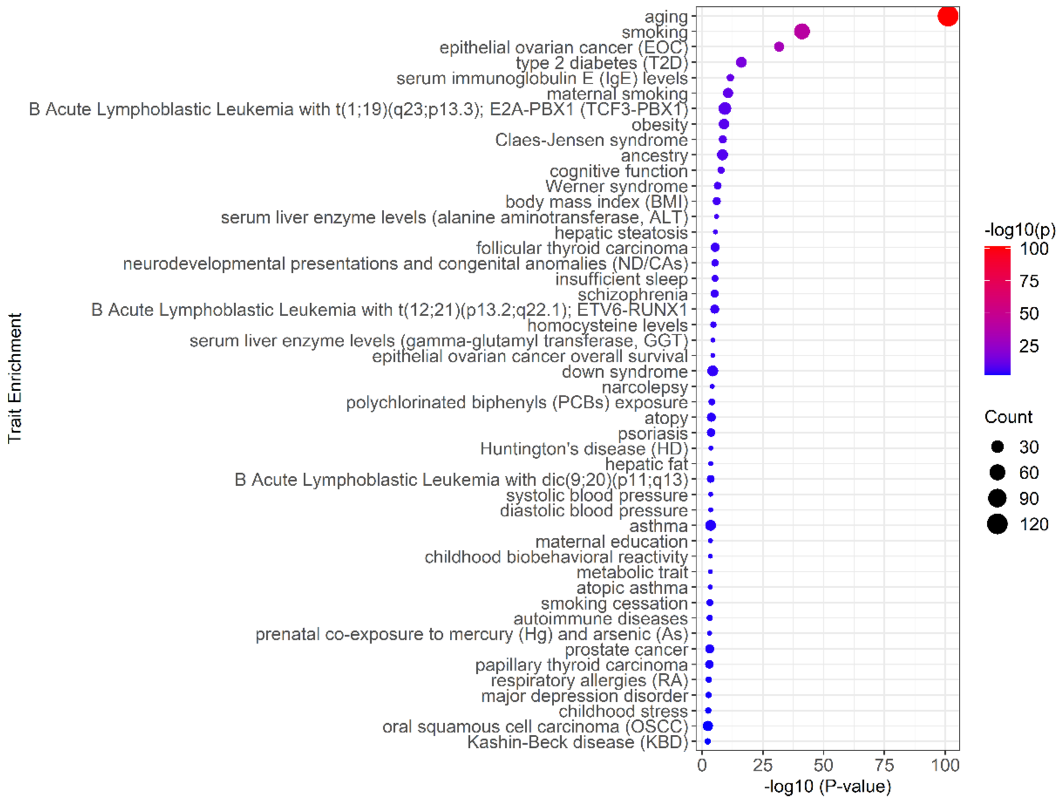

**Fig. S1. Trait enrichment analysis. (a)** Trait enrichment using 353 CpGs selected in HorvathDNAmAge. **(b)** 71 CpGs selected in HannumDNAmAge. **(c)** 513 CpGs selected in PhenoDNAmAge.

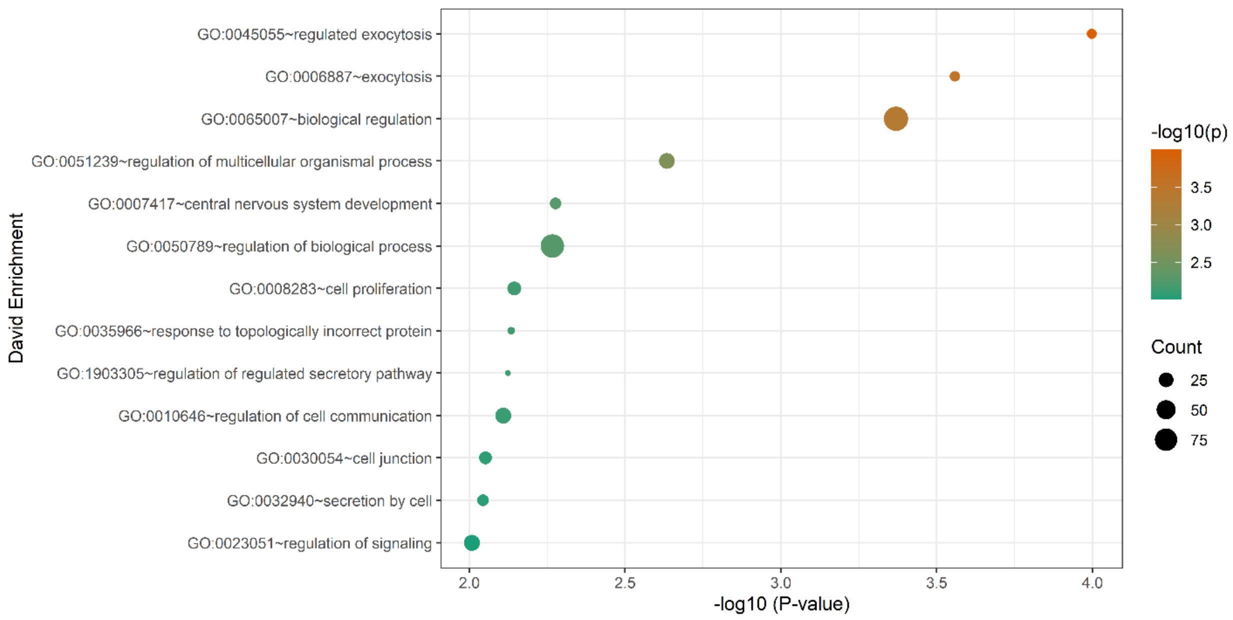

**Fig. S2. Gene set enrichment analysis of novel monocyte DNA methylation age comprising 186 CpGs selected in CD14+ monocyte using DAVID.** DAVID is the web-accessible, gene annotation term-based Database for Annotation, Visualization, and Integrated Discovery.

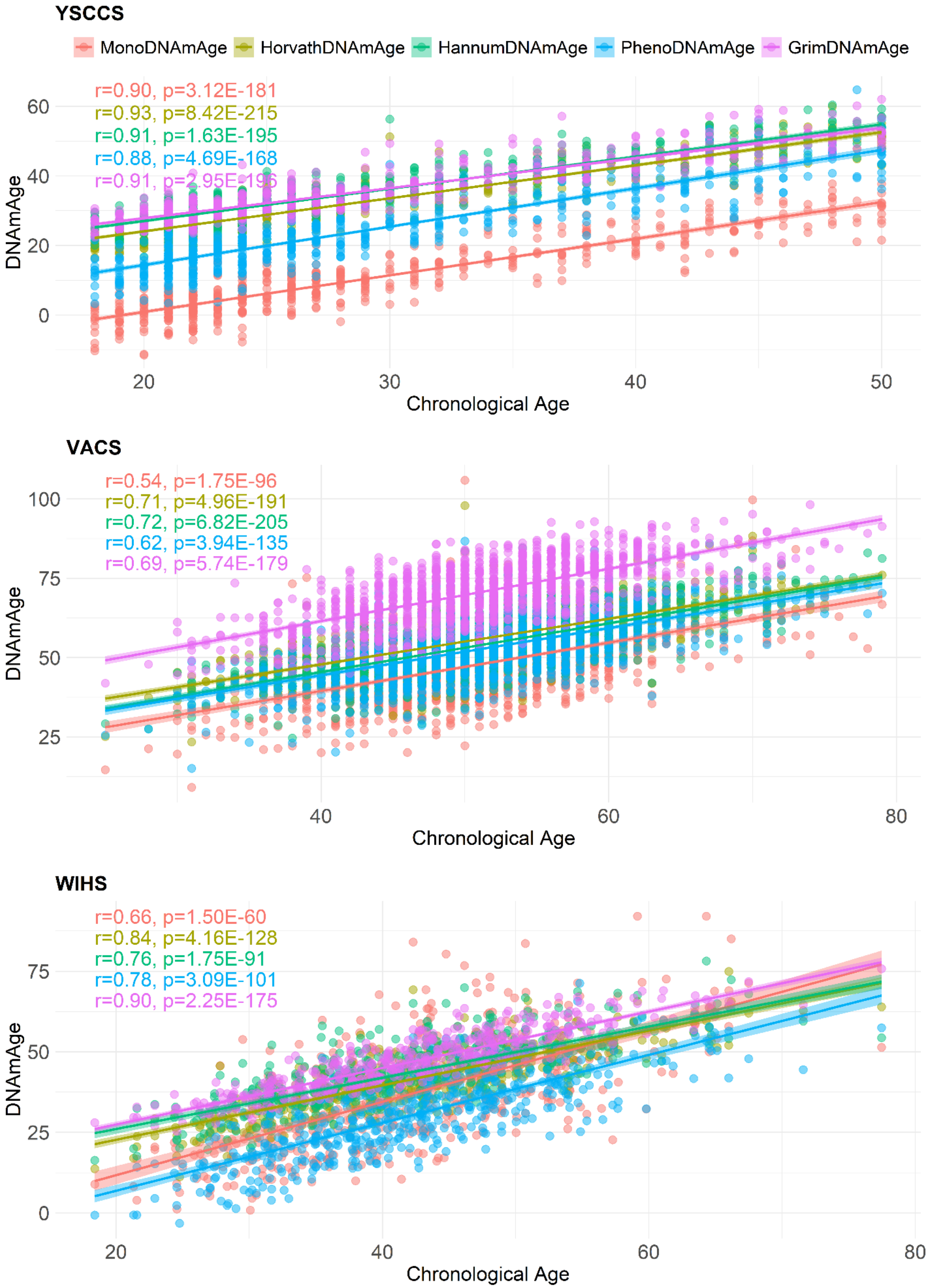
YSCCS
MonoDNAmAge
HorvathDNAmAge
HannumDNAmAge
PhenoDNAmAge
GrimDNAmAge
r=0.90, p=3.12E-181
r=0.93, p=8.42E-215
r=0.91, p=1.63E-195
r=0.88, p=4.69E-168
r=0.91, p=2.95E-196
DNAmAge
Chronological Age
VACS
r=0.54, p=1.75E-96
r=0.71, p=4.96E-191
r=0.72, p=6.82E-205
r=0.62, p=3.94E-135
r=0.69, p=5.74E-179
DNAmAge
Chronological Age
WIHS
r=0.66, p=1.50E-60
r=0.84, p=4.16E-128
r=0.76, p=1.75E-91
r=0.78, p=3.09E-101
r=0.90, p=2.25E-175
DNAmAge
Chronological Age

**Fig. S3. Correlation between five epigenetic clocks and chronological age in three cohorts.** The three cohorts are Yale Stress Center Cohort Study (YSCCS), Veterans Aging Cohort Study (VACS), and Women's Interagency HIV Study (WIHS). The novel clock developed in this study is based on 186 CpGs selected in CD14+ monocyte (MonoDNAmAge); Horvath's 2013 clock based on 353 CpGs selected in multi-tissues (HorvathDNAmAge); Hannum's clock is based on 71 CpGs in leukocytes and was made for adult blood samples (HannumDNAmAge); Levine's DNAm PhenoAge clock employed 513 CpGs selected based on phenotypic age (PhenoDNAmAge); Lu's GrimDNAmAge clock is a linear combination of chronological age, sex, and DNA methylation-based surrogate biomarkers for seven plasma proteins and smoking pack-years (GrimDNAmAge) using 1030 CpGs.

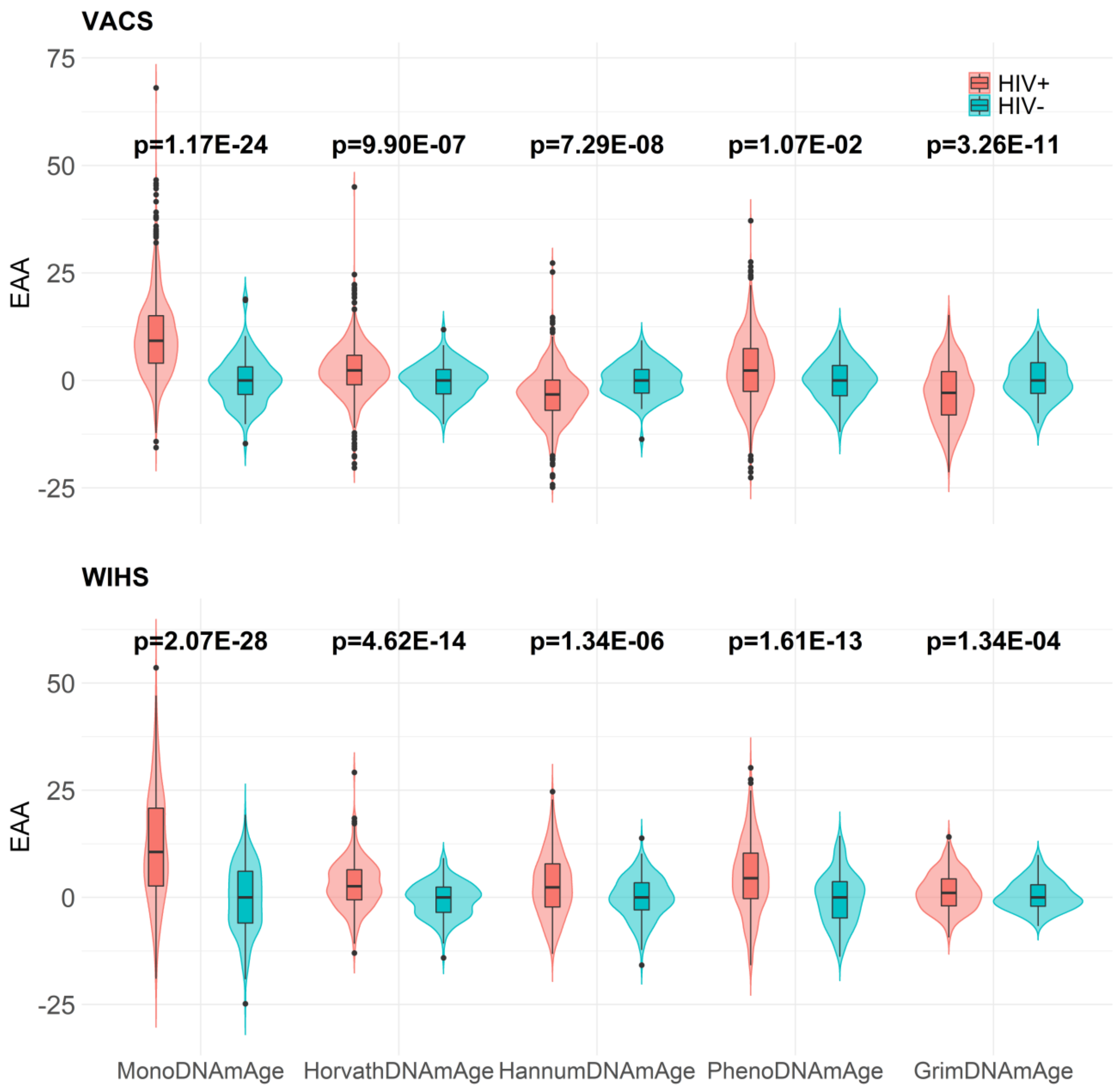

**Fig. S4. Violin plots showing significant differences in EAA between HIV+ and HIV-.** The plots show the Epigenetic Age Acceleration (EAA, the residuals of regressing DNA methylation age on chronological age) between HIV-positive (HIV+) participants and HIV-negative (HIV-) participants in the Veterans Aging Cohort Study (VACS) and the Women's Interagency HIV Study (WIHS).

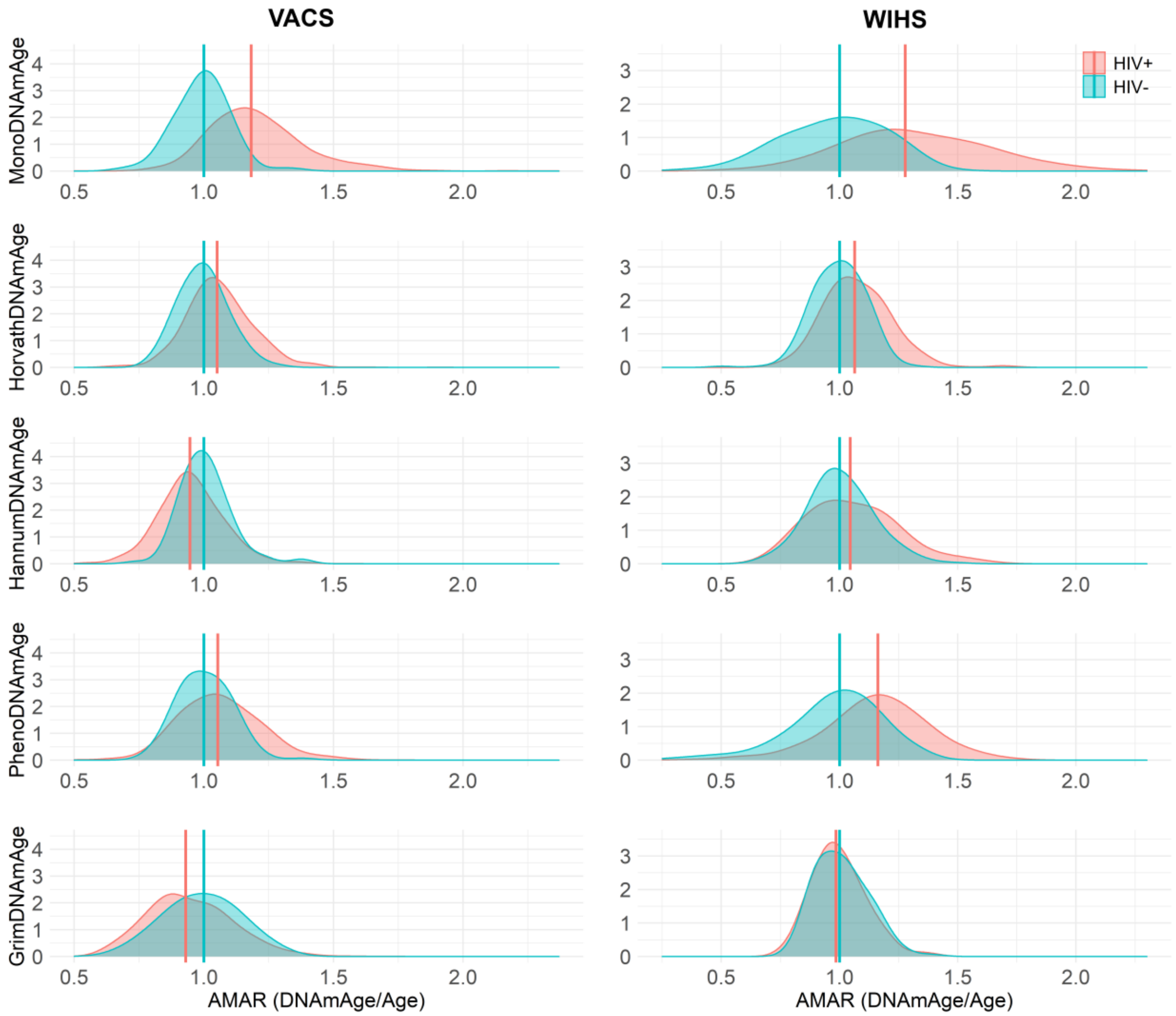

**Fig. S5. Density plots for AMAR between HIV+ and HIV-.** The plots show the Apparent Methylation Age Rate (AMAR, the ratio of DNA methylated age to chronological age) between HIV-positive (HIV+) participants and HIV-negative (HIV-) participants in the Veterans Aging Cohort Study (VACS) and the Women's Interagency HIV Study (WIHS).

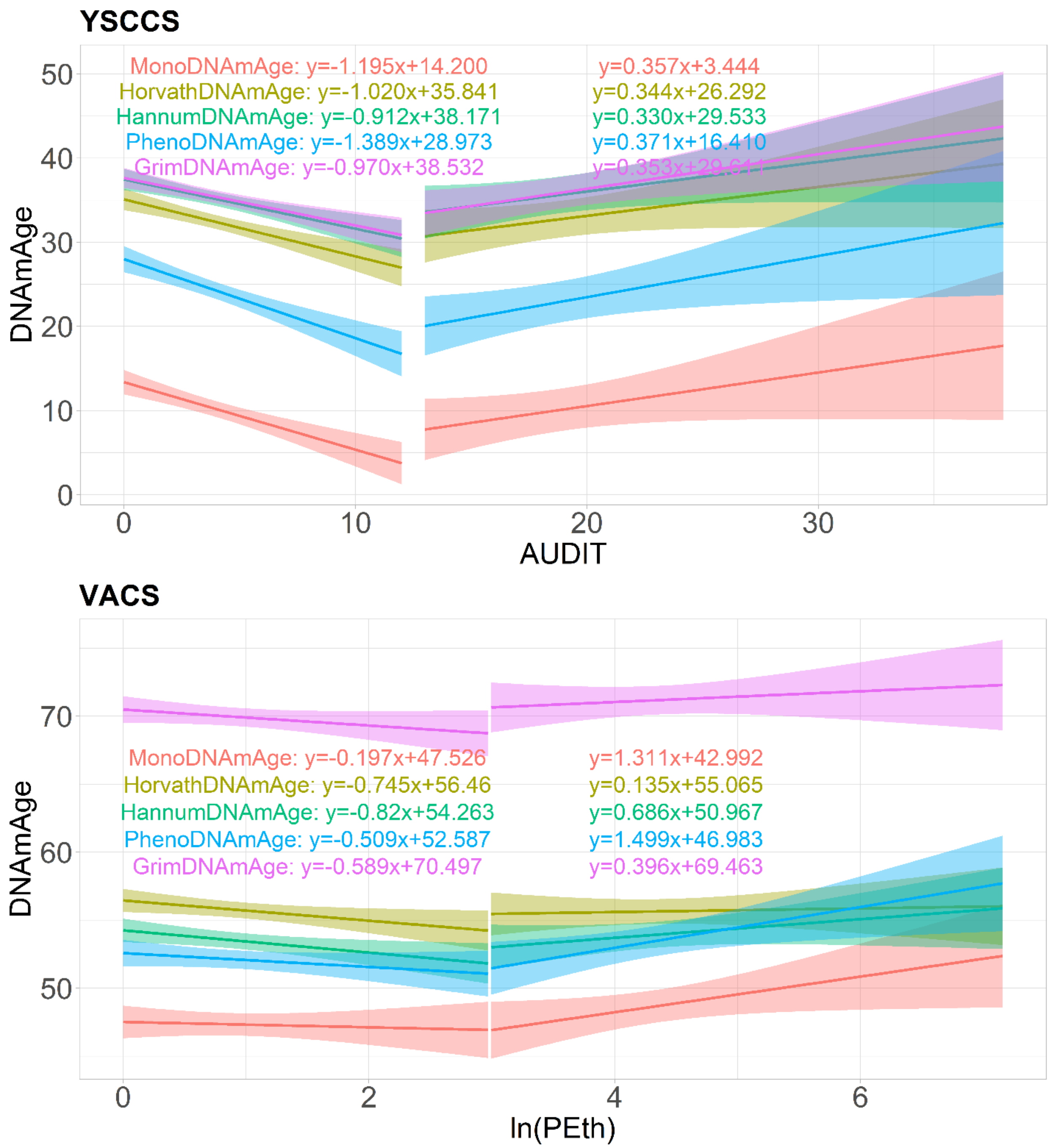

**Fig. S6. Linear regression of DNA methylation age on alcohol consumption.** The plots show a linear association between DNA methylation age (DNAmAge) and light to moderate, and heavy alcohol use separately in the Yale Stress Center Cohort Study (YSCCS) and the Veterans Aging Cohort Study (VACS).

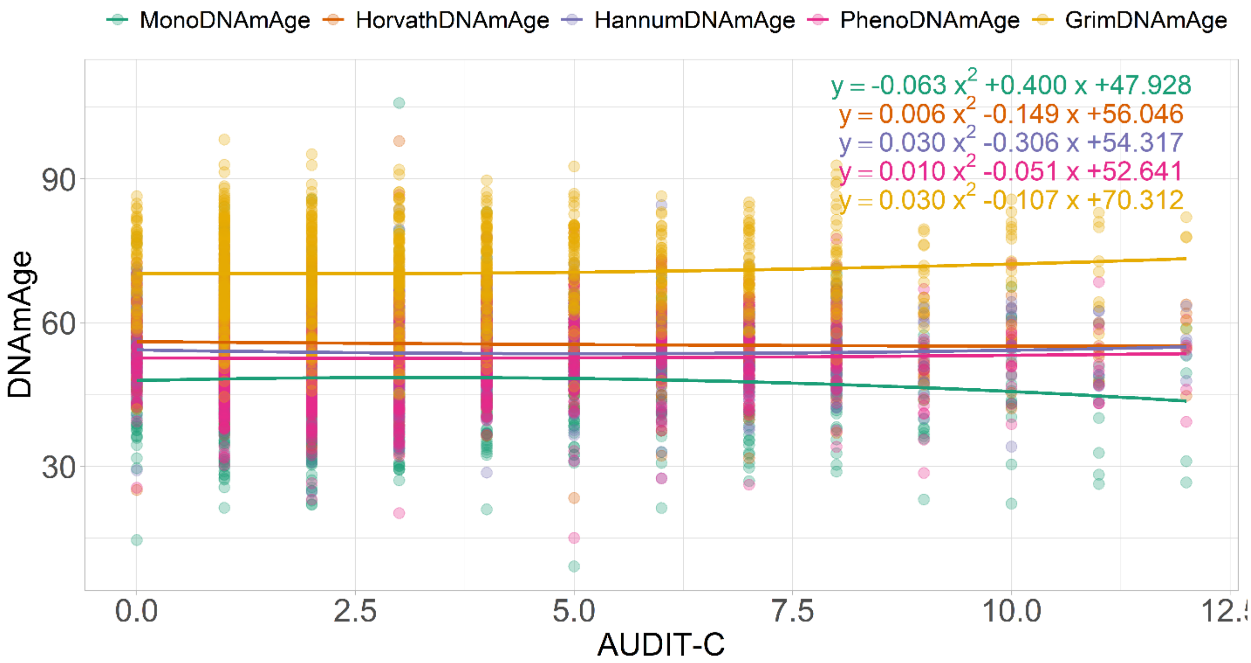

**Fig. S7. Quadratic regression of DNA methylation age on AUDIT-C score.**

Quadratic regression of DNA methylation age (DNAmAge) estimated by five clocks on self-reported Alcohol Use Disorders Identification Test-Consumption (AUDIT-C, first 3 items of AUDIT) score in the Veteran Age Cohort Study (VACS).

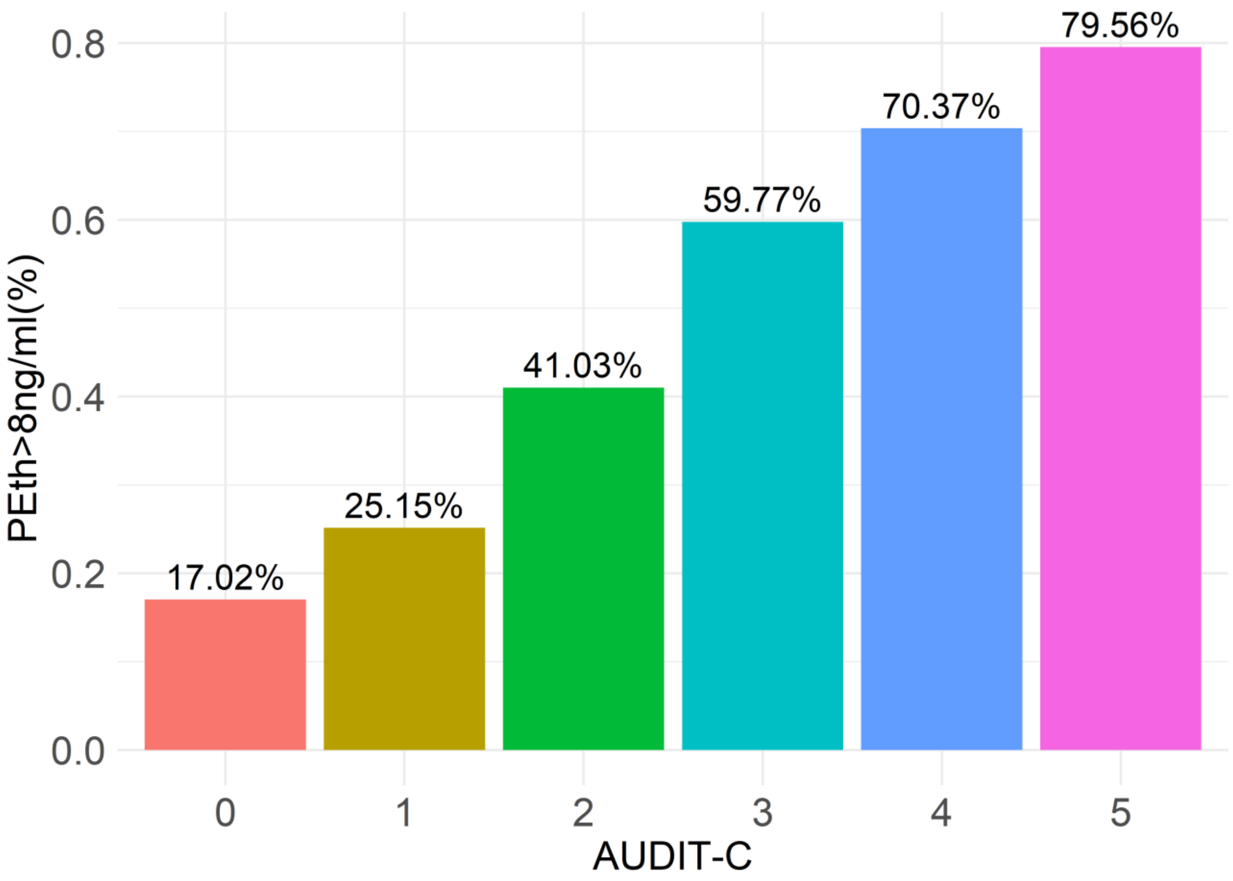

**Fig. S8. Percent with PEth>8 ng/ml by AUDIT-C.** The histogram shows the relationship between objectively measured alcohol use (Phosphatidylethanol (PEth) in the Veterans Aging Cohort Study (VACS)) and self-reported alcohol consumption (Alcohol Use Disorder Identification Test-Consumption (AUDIT-C)).
